# Data quantity is more important than its spatial bias for predictive species distribution modelling

**DOI:** 10.1101/2020.05.24.113415

**Authors:** Willson Gaul, Dinara Sadykova, Hannah J. White, Lupe León-Sánchez, Paul Caplat, Mark C. Emmerson, Jon M. Yearsley

**Affiliations:** School of Biology & Environmental Sciences, Earth Institute, University College Dublin, Dublin, Ireland; School of Biological Sciences, Queen’s University Belfast, Belfast, UK

## Abstract

Biological records are often the data of choice for training predictive species distribution models (SDMs), but spatial sampling bias is pervasive in biological records data at multiple spatial scales and is thought to impair the performance of SDMs. We simulated presences and absences of virtual species as well as the process of recording these species to evaluate the effect on species distribution model prediction performance of 1) spatial bias in training data, 2) sample size (the average number of observations per species), and 3) the choice of species distribution modelling method. Our approach is novel in quantifying and applying real-world spatial sampling biases to simulated data. Spatial bias in training data decreased species distribution model prediction performance, but only when the bias was relatively strong. Sample size and the choice of modelling method were more important than spatial bias in determining the prediction performance of species distribution models.

## 1 INTRODUCTION

Biological records data (“what, where, when” records of species identity, location, and date of observation) often contain large amounts of data about species occurrences over large spatial areas (Isaac & Pocock, 2015). Knowing the geographic areas occupied by species is important for practical and fundamental research in a variety of disciplines. Epidemiologists use maps of predicted wildlife distributions to identify areas at high risk for wildlife-human transmission (Deka & Morshed, 2018; Redding et al., 2019). Land managers can use knowledge of species distributions in spatial planning to minimize impacts on wildlife of new infrastructure (Dyer et al. 2017; Newson et al., 2017). Because complete population censuses are not available for most species, species distribution models (SDMs) are often used to predict distributions of species using relatively sparse observations of species. Species observation data used to train SDMs must represent the study area, but when studies focus on scales of thousands (or tens- or hundreds of thousands) of square kilometers, it is difficult and often expensive to collect adequate data across the entire study extent. Spatially random or stratified sampling of species across large spatial areas is possible, and such surveys exist for some taxa including butterflies and birds (Uzarski et al., 2017), but such data are uncommon for most taxonomic groups (Isaac, van Strien, August, de Zeeuw, & Roy, 2014). More commonly, data are either spatially extensive but collected opportunistically (Amano, Lamming, & Sutherland, 2016), or are collected according to structured study designs but are more spatially limited.

Collecting biological records data is relatively cheap compared to collecting data directly as part of a research project (or at least the costs of collecting biological records are borne in large part by individual observers rather than by data analysts) (Carvell et al., 2016). However, there is an associated challenge because the analyst lacks control over where, when, and how data were collected. Many biases have been documented in biological records data, including temporal, spatial, and taxonomic biases (Boakes et al., 2010). Spatial sampling bias, in which some areas are sampled preferentially, is particularly pervasive at all scales and across taxonomic groups (Amano & Sutherland, 2013; Oliveira et al., 2016). Despite these biases, biological records are often used in species distribution modelling, either because no other data exists at the spatial scale of interest, or because the modeler expects biological records to be more informative than data from more explicitly designed but smaller sampling schemes. Given the ubiquitous presence of spatial sampling bias in biological records data, it is important to know whether spatial bias in training data impedes the ability of SDMs to correctly model species distributions. Data collection efforts often face a practical trade-off between maximizing the overall quantity and the spatial evenness of new records. It would thus be useful to know whether the value of biological records for SDMs can best be improved by increasing the spatial evenness of recording (perhaps at the cost of the overall amount of new data that is added), or by increasing the overall amount of recording (even if new records are spatially biased).

Spatial sampling bias in biological records has similarities with sampling biases that have been investigated in other settings. The field of econometrics uses the term “sample selection bias” to refer to non-random sampling and has developed theory about when sampling bias is likely to bias analyses (Wooldridge, 2009). A key consideration in econometrics’ evaluations of sample selection bias is determining whether the inclusion of data in the sample depends on predictor variables that are included in the model (“exogenous” sample selection), or depends on the value of the response variable (“endogenous” sample selection), or both (Wooldridge, 2009). In ecology, Nakagawa (2015) similarly provides guidelines for assessing missing data in terms of whether data is missing randomly or systematically with respect to other variables (see also Gelman & Hill, 2006). In a machine learning context, Fan, Davidson, Zadrozny, & Yu (2005) investigated the effect on predictive models of sample selection bias in which sampling is associated with predictor variables - “exogenous sample selection” in the terms of Wooldridge (2009) and “missing at random” in the terms of Nakagawa (2015) - and determined that most predictive models could be sensitive or insensitive to sampling bias depending on particular details of the dataset.

Biological records may have been collected with spatial sampling biases that are exogenous, endogenous, or both, and datasets may contain a mix of records collected with different types of bias. For example, when sampling intensity depends on proximity to roads (Oliveira et al., 2016), the sampling bias is exogenous because records arise from biased sampling that depends on an aspect of environmental space that can be included in models as a predictor variable. However, when a birder, for example, submits a record of an unusual bird from a location where they would not otherwise have submitted records, the bias is endogenous because the sampling location depends on the value of the response variable (species presence). In reality, the observer might have seen the unusual bird while driving along a road, so the sampling location depends on both the response variable (the presence of the bird) and predictor variables (proximity to the road). Most sampling biases occur on a continuum and are not unequivocally categorizable using any existing scheme (Nakagawa, 2015), making it difficult to describe exactly the biases in data or predict their effect on model performance.

Studies testing the impact of spatially biased training data on predictive SDMs have shown mixed results. Multiple studies using a pseudo-absence (or “presence/background”) approach with presence-only biological records have found that spatial bias in the data used to train SDMs decreases model prediction performance (Phillips et al., 2009; Barbet-Massin, Jiguet, Albert, & Thuiller, 2012; Stolar & Nielsen, 2015). However, it is not clear whether the effect of the spatial bias in those cases is due to the bias in the original data or the relative difference in bias between the original data and pseudo-absences. In fact, Phillips et al. (2009) found that spatial bias in the presence records strongly reduced model performance when using a pseudo-absence approach but not when using a presence-absence approach. Some SDM methods tested by Barbet-Massin et al. (2012) appeared relatively unaffected by spatial sampling bias, while generalized linear models (GLMs) and generalized additive models (GAMs) appeared to be more strongly affected. Classification trees were sensitive to spatially biased training data in a study of lichen distributions (Edwards, Cutler, Zimmerman, Geiser, & Moisen, 2006). Thibaud, Petitpierre, Broennimann, Davison, & Guisan (2014) found that the effect of spatial sampling bias on SDM prediction performance depended on the SDM modelling method, and that the effect of spatial sampling bias was smaller than the effect of other factors, including sample size and choice of modelling method. Warton, Renner, & Ramp (2013) provided a method for correcting for spatially biased data when building SDMs, but found that the resulting improvement in model predictive performance was small. Because there is no clear guidance about when spatial bias in training data will or will not affect model predictions, tests of the observed effect of spatial biases common in biological records are important for determining whether those biases are likely to be problematic in practice.

The effect of spatial sampling bias on model predictions can be studied using either real or simulated data (Zurell et al., 2010). Using real data has the advantage that the biases in the data are, well, real. The spatial pattern, intensity, and correlation of sampling bias with environmental space are exactly of the type that analyses of real data must cope with. However, using real data has two disadvantages. First, the truth about the outcome being modeled (species presence or absence) is not completely known in the real world, making it impossible to evaluate how well models represent the truth. Second, biases in real data are not limited to the biases under study – a study investigating the effect of exogenous spatial sampling bias will be unable to exclude from a real dataset records generated by endogenously biased sampling that depends on the values of the outcome variable. Simulation studies avoid both these problems. Because the investigator specifies the underlying pattern that is subsequently modeled, the truth is known exactly (even when realized instances of the simulation are generated with some stochasticity). The investigator also has direct control over which biases are introduced into a simulated dataset, and therefore can be more confident that any observed effects on predictions are due to the biases under investigation.

Spatial sampling bias can be introduced into either simulated or real data. This can be done using a parametric function that describes the bias (Isaac et al., 2014; Stolar & Nielsen, 2015; Thibaud et al., 2014) or by following a simplified ad-hoc rule (e.g. splitting the study region into distinct areas that are sampled with different intensities) (Phillips et al., 2009). However, these approaches may not adequately test the effect of spatial bias if the biases found in real biological records do not follow parametric functions or are more severe than artificial parametric or ad-hoc biases. We used observed sampling patterns from Irish biological records to sample simulated species distributions using realistic spatially biased sampling.

We used a virtual ecologist approach (Zurell et al., 2010) applied at the scale of Ireland to investigate the effect on the predictive performance of SDMs of 1) spatial sampling bias, 2) sample size (the average number of records per species), and 3) choice of SDM method. Our method for introducing sampling bias preserves real-world spatial patterns of sampling bias at multiple scales - not only are some individual locations more heavily sampled than others, but heavily sampled locations are arranged in the landscape non-randomly in relation to each other and in relation to the landscape itself (i.e. some habitats are better sampled than others). We quantified the spatial sampling biases used in our study to enable comparison with biases in other datasets. Our approach is novel in applying real-world spatial sampling biases to simulated data.

## 2 METHODS

We assessed the ability of species distribution models to predict “virtual species” distributions (Leroy, Meynard, Bellard, & Courchamp, 2016; Zurell et al., 2010) when the models were trained with datasets with a range of spatial sampling biases and sample sizes. Virtual species distributions were produced by defining the responses of virtual species to environmental predictor variables (Table 1). Occurrence maps for virtual species were based on the actual values of the environmental predictor variables in 840 10 km x 10 km grid squares in Ireland (total area of study extent = 84,000 km^2^). We generated “virtual biological records” by sampling the community of virtual species in each grid square using sampling patterns taken from Irish biological records data.

**Table 1.** Environmental predictor variables used to define and model the distribution of virtual species in Ireland. Moran’s I values indicate the spatial clustering of values for each variable, where a value of one indicates strong spatial clustering of variable values, zero indicates random spatial arrangement of values, and negative one indicates strongly dispersed spatial arrangement of values. Details of data sources are in Section 2.1.

| Variable | Description | Data Source | Moran's I |
| --- | --- | --- | --- |
| annual minimum temperature (degrees C) | 2% quantile of annual temperatures in each grid cell averaged over the years 1995-2016 | E-OBS | 0.84 |
| annual maximum temperature (degrees C) | 98% quantile of annual temperatures in each grid cell averaged over the years 1995-2016 | E-OBS | 0.83 |
| annual precipitation (mm) | Average total annual precipitation in each grid cell over the years 1995-2016 (excluding 2010-2012) | E-OBS | 0.82 |
| average daily sea level atmospheric pressure (hecto Pascals) | Average daily sea level atmospheric pressure over the years 1995-2016 | E-OBS | 0.86 |
| agricultural areas | Proportion of each grid cell classified as agricultural areas | CORINE Land Cover Database | 0.53 |
| artificial surfaces | Proportion of each grid cell classified as artificial surfaces | CORINE Land Cover Database | 0.44 |
| forest and semi-natural areas | Proportion of each grid cell classified as forest and semi-natural areas | CORINE Land Cover Database | 0.41 |
| water bodies | Proportion of each grid cell classified as water bodies | CORINE Land Cover Database | 0.35 |
| wetlands | Proportion of each grid cell classified as wetlands | CORINE Land Cover Database | 0.55 |
| elevation | Average elevation in each grid cell | ETOPO1 | 0.29 |

### 2.1 Environmental predictor variables

We chose environmental predictor variables with a range of spatial patterns and scales of spatial auto-correlation (Table 1, Fig. S1). Because our species were simulated, predictor variables did not need to have biological relevance - by definition, the variables used to create the range of each virtual species were relevant to that species. The variety of spatial patterns in our predictor variables ensured that our virtual species distributions were determined by variables with a variety of spatial patterns, as is the case for real biological species. We used climate variables (which show relatively strong spatial clustering, Table 1) from the E-OBS European Climate Assessment and Dataset EU project (Haylock et al., 2008; van den Besselaar, Haylock, van der Schrier, & Klein Tank, 2011; http://www.ecad.eu/download/ensembles/downloadchunks.php). We calculated the proportion of each grid square covered by different land cover variables (which show less spatial clustering than climate variables, Table 1) from the CORINE Land Cover database (CORINE, 2012). We calculated the average elevation within each grid square by interpolation using ordinary kriging from the ETOPO1 Global Relief Model (Amante & Eakins, 2009; https://www.ngdc.noaa.gov/mgg/global/relief/ETOPO1/data/ice_surface/grid_registered/netcdf/ [accessed 8 May 2019]).

Spatial data were prepared using the ‘sf’, ‘sp’, ‘raster’, ‘fasterize’, ‘rgdal’, ‘gstat’, and ‘tidyverse’ packages in R version 3.6 (Bivand, Keitt, & Rowlingson, 2018; Gräler, Pebesma, & Heuvelink, 2016; Hijmans 2018; Pebesma, 2018; R Core Team, 2018; Ross, 2018; Wickham, 2017).

### 2.2 Species occurrence data

We downloaded observations of species across the island of Ireland for the years 1970 to 2014 from the British Bryological Society for bryophytes (accessed through NBN Atlas website, https://nbnatlas.org [downloaded 24 August 2017]) and from the Irish National Biodiversity Data Centre (NBDC) for moths, dragonflies, butterflies, and birds (http://www.biodiversityireland.ie/ [downloaded 6 October 2017]). The data contained presence-only records of species, with the date and location of the observation, an anonymized observer identifier, and a taxonomic group label that indicated species commonly sampled together. The taxonomic group label often corresponded to order (e.g. odonata), but sometimes represented a class (e.g. Aves) or other categorization that better grouped species according to sampling techniques. Locations of records were provided as either 1 km^2^ or 100 km^2^ (10 km x 10 km) grid squares, but we used 10 km x 10 km grid squares in all analyses in order to retain the majority of the data. Within each taxonomic group, we grouped records into sampling events, where a sampling event was defined as all records with an identical combination of recording date, location, and observer.

### 2.3 Spatial sampling patterns in Irish species occurrence data

For each taxonomic group, we quantified sampling effort in each grid square as the proportion of all records coming from the grid square. We used grid squares along the coast even though these cells contain less terrestrial habitat than inland grid squares. We measured the spatial evenness of sampling effort among locations by using Simpson evenness (Magurran & McGill, 2011) to compare the number of observation records in grid squares.

### 2.4 Data simulation

#### 2.4.1 Simulating species distributions

We simulated and sampled virtual species distributions using the ‘virtualspecies’ package (Leroy et al., 2016) in R. The probability of occurrence of each virtual species *i* in each grid square *j* was a logistic function of two variables and their quadratic terms:

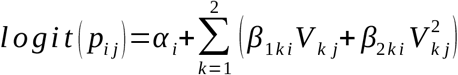

where *p_ij_* is the probability that virtual species *i* occurs in grid square *j, V_kj_* indicates the value of the *k*^th^ predictor variable in grid cell *j*, and the *α* and *β* terms are the species-specific coefficients defining the response of the virtual species to the environment. The predictor variables were derived by randomly selecting, for each virtual species, seven of the ten environmental variables to use as drivers of occurrence (only seven of the ten variables were used for each species so that not all species responded to all the same environmental variables). Selected environmental variables were centered, scaled, and summarized using principal components analysis with the ‘ade4’ R package (Dray & Dufour, 2007). The first two principal components were used to determine the distribution of the species, rather than using the seven original environmental variables, to avoid producing virtual species with optimal niches in conditions that do not exist (e.g. a virtual species with an occurrence optimum at warm temperature and high elevation) (Leroy et al., 2016). Coefficients specifying virtual species’ responses were chosen such that the theoretical prevalence of each virtual species (the sum of the probabilities of presence in each grid square divided by the number of grid squares) was greater than 0.01, equivalent to the virtual species occurring in at least eight of the 840 grid squares in our study extent.

#### 2.4.2 Realized species communities

A single realized distribution of each virtual species *i* was created by randomly generating a “presence” (1) or “absence” (0) for each grid square *j* by drawing a value from a binomial distribution with probability *p_ij_*. We simulated two different types of virtual species communities, a small community containing 34 virtual species (the number of recorded odonata species in Ireland) and a large community containing 1268 virtual species (the number of recorded bryophyte species in Ireland). Results were qualitatively similar for the large- and small-community simulations after fitting two of the SDM methods (GLMs and inverse distance-weighted interpolation). We therefore tested the third SDM method, boosted regression trees, only on the large-community simulation. Below we refer to the large community simulation except where explicitly stated. For small community simulation results see supplementary materials (S2).

#### 2.4.3 Simulating sampling with spatial bias

Virtual biological records data were generated by sampling the realized species communities in “sampling events” at different locations to produce spatially explicit species checklists (Fig. S3). Spatial sampling locations were chosen based on spatial sampling patterns from three Irish biological records datasets with different spatial sampling biases: birds (low spatial sampling bias), butterflies (median spatial sampling bias), and moths (severe spatial sampling bias). This gave four spatial sampling “templates”, including the case of no spatial sampling bias (Fig. 1).

**Fig. 1.**
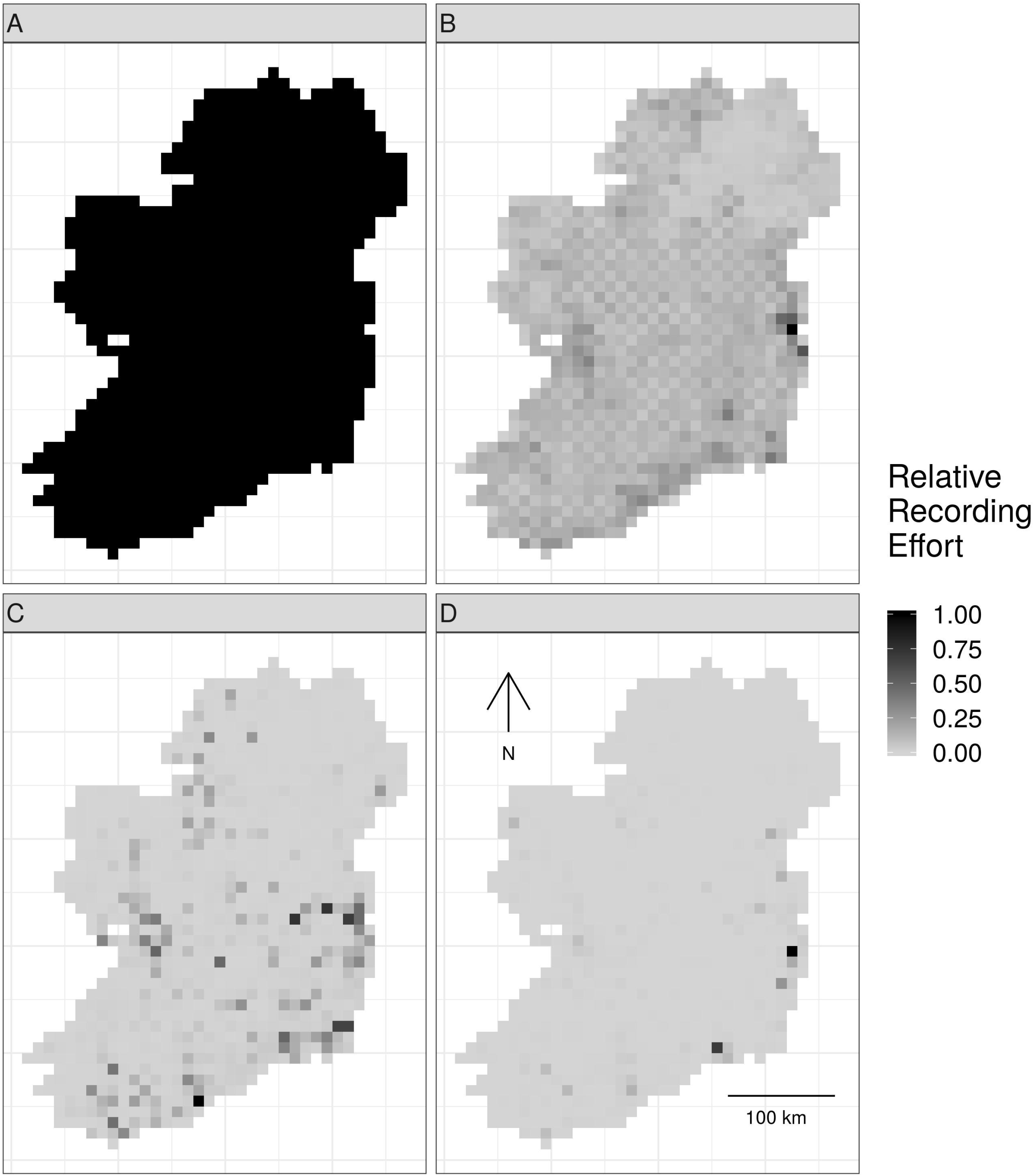
Spatial sampling patterns from Irish biological records. Spatial sampling patterns from Irish biological records were used as templates to create virtual species records data with varying amounts of spatial bias. Darker shades indicate higher relative probability of sampling from a grid square compared to other grid squares within in the same template; overall sampling effort is the same for each panel (A) through (E). The most heavily sampled grid square in each spatial bias template has a relative recording effort of one, while a grid square with half as many records as the most heavily sampled square has a relative recording effort of 0.5. Spatial sampling patterns derived from datasets for different taxonomic groups were: (A) no bias (even probability of sampling from every grid square), (B) low bias (based on bird data), (C) median bias (based on butterflies), and (D) severe bias (based on moths).

To make sampling patterns comparable between datasets with different sample sizes, we calculated sampling weights for each grid square in each empirical dataset by counting the number of records in each grid square and dividing by the maximum number of records in any grid square. This produced a relative sampling weight for each grid square, where the most heavily sampled cell had a weight of one and other cells had weights below one (Fig. 1).

We tested six different sample sizes, defined as the mean number of records per species (number of records per species = 2, 5, 10, 50, 100, and 200).

To generate virtual biological records from the virtual species communities, we randomly selected a grid square, using selection probabilities from one of the four spatial-bias templates. Within each grid square that was selected for sampling, we 1) generated a list of virtual species that were present in the grid square; 2) defined the probability of observing each of the present species based on the species’ prevalence in the entire study extent (so that common species had a higher probability of being recorded when present), and 3) drew observations with replacement from the list of present species. The number of records to generate during a sampling event (i.e. the checklist length) was drawn randomly with replacement from the sampling event checklist lengths from real bryophyte data (for the large community simulation) or dragonfly data (for the small community simulation). We continued this sampling process until we had accumulated the desired number of records.

### 2.5 Species distribution modeling

We tested three different SDM modeling techniques: generalized linear models (GLMs) (Hosmer & Lemeshow, 2000), boosted regression trees (Elith, Leathwick, & Hastie, 2008; Friedman, 2001), and inverse distance-weighted interpolation (Cressie, 1991). These represent distinct types of methods used for SDMs, including linear (GLM) and machine learning (boosted regression tree) methods, and a spatial interpolation method (inverse distance-weighted interpolation) that does not include information from environmental covariates. For all methods, the modeled quantity was the probability of the focal virtual species being recorded on a checklist. We modeled each species individually as a function of five environmental predictor variables, chosen from the ten possible predictor variables listed in Table 1. Using only five of the ten possible predictor variables simulated a real-world situation in which the factors that influence species distributions are not entirely known. We treated the list of records from each sampling event as a complete record of that sampling event, and treated the absence of species from a sampling event checklist as non-detection data for those species (Fig. S3, Kéry et al., 2010). Thus, we explicitly used a detection/non-detection rather than a presence-only modeling framework. Many species distribution modelling techniques commonly used with presence-only data require the generation of artificial “pseudo-absences” in order to fit models (Barbet-Massin et al. 2012). However, the spatial bias of pseudo-absences should match the spatial bias of presence data, which can be difficult to achieve, especially when spatial biases are difficult to model. We avoided the use of pseudo-absences by analyzing checklists of species, on which every species is either detected or not detected (Johnston et al. 2020, Kéry et al. 2010). Using non-detection data inferred from records of other similar species provides clarity about what is being modeled (i.e. the probability of a species being recorded on a checklist, not the probability of occurrence) and ensures that the sampling biases are the same for detections and nondetections, which may reduce the effect of sampling bias (Barbet-Massin et al. 2012, Johnston et al. 2020, Phillips et al. 2009).

We modeled 110 randomly selected virtual species from the 1268 virtual species in the large community simulation. The number of virtual species modeled was a compromise between high replication and computation limitations, but testing 110 virtual species should provide enough replication for robust conclusions. We fitted each type of SDM once to each combination of virtual species, sample size, and spatial sampling bias. Thus, the sample size for our study - the number of SDM prediction performance values that we used to assess the effects of spatial sampling bias, sample size, and SDM method - was 110 prediction performance values for each combination of SDM method, sample size, and spatial sampling bias (one prediction performance value for each of the 110 selected virtual species). Replication in our study came not from repeatedly fitting models to different randomly generated sets of presences and absences of the same virtual species, but rather from fitting each model once to data for many different virtual species, all generated using parameters randomly drawn from the same distributions. However, the same 110 virtual species were used for each combination of SDM method, spatial sampling bias, and sample size, ensuring that all comparisons were based on the same underlying task (i.e. modelling the same true species distributions).

Models were trained and evaluated using five-fold spatial block cross-validation (Roberts et al., 2017) that partitioned the study extent into spatial blocks of 100 km x 100 km and allocated each block to one of five cross-validation partitions. Models were trained five times, each time leaving out data from one of the five partitions. We only attempted to fit models if there were more than five positive detections in the training data (i.e. within the four training folds during crossvalidation), because we did not expect any of the SDM methods we tested to be able to produce meaningful models when there were fewer than six detections of the focal species. Prediction performance of models was evaluated using the true simulated species presence or absence in each grid cell not included in the spatial extent of the training partitions (Fig. 2). Thus, evaluation data was spatially even and the number of evaluation points stayed constant even as the sample size and spatial bias of training data changed (Fig. 2). Prediction performance was evaluated using the area under the receiver operating characteristic curve (AUC) (Hosmer & Lemeshow, 2000) to measure models’ ability to accurately distinguish presences and absences, and root mean squared error (RMSE) to compare predicted probabilities of species being recorded during a sampling event to the true probability of occurrence defined by the simulation.

**Fig. 2.**
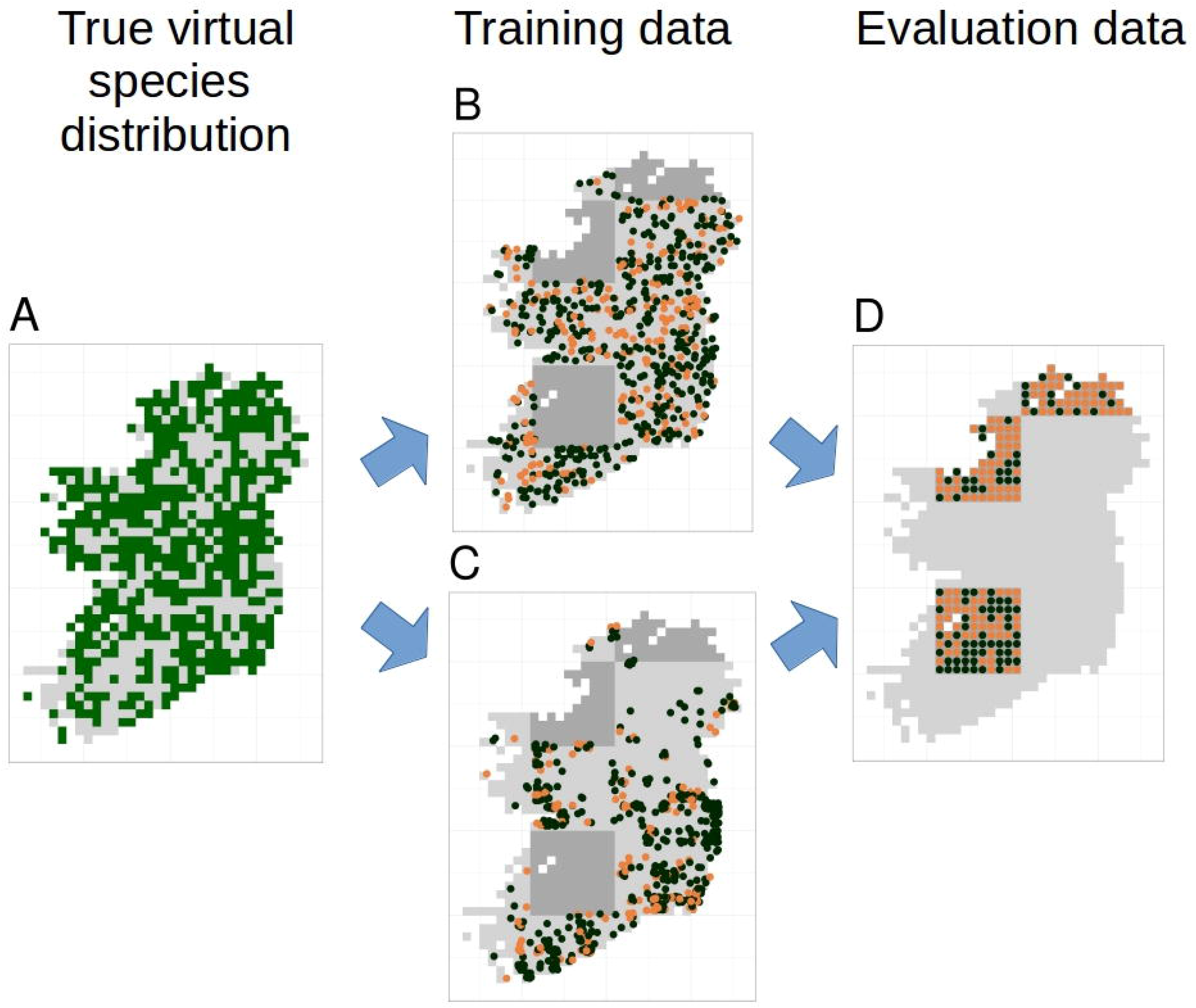
Species distribution model training and testing process for a single cross-validation fold. The true virtual species distribution (A, presences shown in dark green, absences in light grey) was sampled to produce virtual biological records with a range of sample sizes and spatial biases, including no bias (B) and median bias (C). Orange points in (B) and (C) show checklists on which the species was recorded, black points show checklists on which the species was not recorded (i.e. non-detection points). Species distribution models were fit using five-fold spatial block cross validation, in which data from about 80% of the spatial area was used to train models (light grey background in B and C). Data from the remaining spatial areas (dark grey background in B and C) was set aside for model evaluation. Model evaluation tested the ability of species distribution models to predict the true presence (orange dots) or absence (black dots) of the species in each grid cell within the evaluation areas (D). Model evaluation therefore used spatially even data with the same number of evaluation points (D) regardless of the sample size and spatial bias of training data (B and C).

For GLMs, we used logistic regression (‘glm’ function) with a binomial error distribution and logit link. Quadratic terms were fitted, but we did not fit interactions between variables. We controlled overfitting by limiting the number of terms in GLMs such that there were at least 10 detections or non-detections (whichever was smaller) in the training data for each non-intercept term in the model. For example, if the training data had 35 detections, we limited the GLM to using only three terms plus an intercept. We tested all possible models from an intercept-only model up to models with the maximum number of terms permitted by our “10 detections per term” rule of thumb. If a quadratic term was included in a model, we also included the 1^st^ degree term. For generating predictions, we used the model that gave the lowest AIC based on the training data.

Boosted regression trees were trained using ‘gbm.step’ in the ‘dismo’ package (Greenwell, Boehmke, & Cunningham, 2018; Hijmans, Phillips, Leathwick, & Elith, 2017). Unlike GLMs, boosted regression trees do not require the modeler to specify interactions between variables, because the trees will discover and model interactions if they are present. The tree complexity specified by the modeler controls the maximum interaction order that the models are permitted to fit, and therefore can be used to prevent overfitting. Elith, Leathwick and Hastie (2008) found relatively little harm in using higher tree complexities, even with small sample sizes, presumably because the models did not fit complex interactions that were not present, even when the model was given freedom to do so. Nevertheless, we tested tree complexities of two and five, to build models that allowed interactions between up to two and up to five variables, respectively. Smaller learning rates are generally preferred because they result in better predictive performance but using smaller learning rates comes at the cost of higher computation and memory requirements (Elith, Leathwick, and Hastie 2008). We therefore used learning rates small enough to grow at least 1000 trees (following Elith, Leathwick, and Hastie 2008), but large enough to keep models below an upper limit of 30,000 trees because of computation time limitations. We used gbm.step to determine the optimal number of trees for each model, based on monitoring the change in 10-fold cross-validated error rate as trees were added to the model (Hijmans, Phillips, Leathwick, & Elith, 2017). We explored whether the upper limit of 30,000 trees affected our conclusions by looking at graphs of the frequency distribution of number of trees used, and graphs of prediction performance as a function of the number of trees. Details of the procedure used to select the tree complexity, learning rate, and number of trees are in the supplementary materials (S2) and in our R code, which is available on GitHub (https://zenodo.org/badge/latestdoi/229083757).

Inverse distance-weighted interpolation was implemented using ‘gstat’ (Gräler et al., 2016; Pebesma, 2004). We tuned parameters of the inverse distance-weighted interpolation model based on prediction error (details in S2 and at https://zenodo.org/badge/latestdoi/229083757).

After models were fitted, we looked for evidence of overfitting and assessed whether the number of positive detections of the focal species in the test dataset affected prediction performance metrics. Details of the graphs used to assess overfitting and the effect of species prevalence on performance metrics are in the supplementary materials (S2). All analyses used R version 3.6.0 (R Core Team, 2020), and code is available on GitHub (https://zenodo.org/badge/latestdoi/229083757).

### 2.6 Analyzing effects of sampling bias and sample size

We modeled the predictive performance (AUC and RMSE) of SDMs as a function of spatial sampling bias, sample size (average number of observations per species), and SDM method. Modelling was done using boosted regression trees (‘gbm.step’ in the ‘dismo’ package) (Greenwell et al., 2018; Hijmans et al., 2017). To assess whether species prevalence (the commonness or rarity of a species in the study extent) and/or the number of detections in the test dataset affected our evaluations of model performance, we graphed AUC and RMSE as a function of species prevalence for all models (Fig. S4), and graphed AUC as a function of the number of detections in the test dataset for each SDM modelling method separately (Fig. S5). Because RMSE showed a strong trend with species prevalence (Fig. S4), we included species prevalence in the boosted regression tree models of RMSE. AUC showed decreasing variability as prevalence increased, but did not show a clear trend that was not associated with the decrease in variability (Fig. S4). AUC did not show any trend with the number of detections in the test dataset (Fig. S4). Because AUC did not seem to be strongly affected by species prevalence or the number of detection in the test data, we did not include species prevalence in our models assessing AUC. Variable importance was assessed based on the reduction in squared error attributed to each variable in boosted regression tree models (Friedman, 2001). We also assessed the effect of spatial sampling bias and sample size of training data on the number of species for which models could be fitted within the computational time and memory constraints of this study (S2).

## 3 RESULTS

Simulated species showed a variety of plausible distribution patterns (Fig. 3) and prevalences (Fig. S6), including species with north/south distribution gradients and distributions that followed geographic features such as the coastline (Fig. 3).

**Fig. 3.**
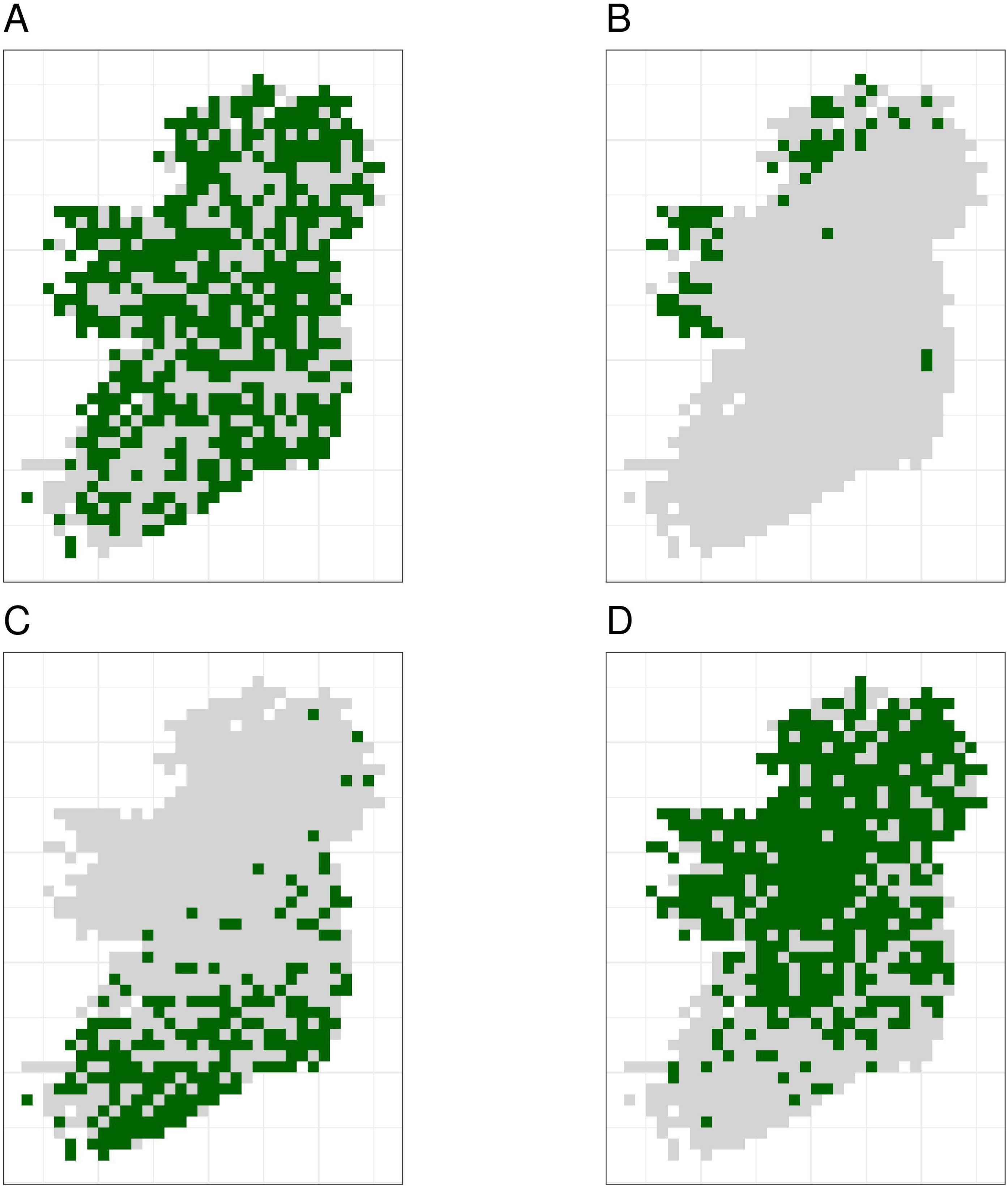
The true distributions of four example simulated species. Simulated species showed a range of plausible distributions with a range of prevalences, including (A) common widespread species, (B) rare species mostly limited to north-western coastal sites, (C) species with a north/south gradient in occurrence, and (D) common species that are absent from southern sites.

Sample size (the mean number of observations per species) was the most important variable for explaining variations in prediction performance of SDMs, followed by the choice of SDM method and spatial sampling bias (Table 2). Simpson evenness values for spatial sampling evenness of the template datasets are in Table 3.

**Table 2.** Importance of sample size, spatial bias, and modelling method for determining predictive performance of species distribution models. Variable importance measures from a boosted regression tree show the relative influence of sample size (average number of records per species), species distribution modeling method, and spatial bias in training data on prediction performance (AUC) of species distribution models. The relative influence for each variable is the reduction in squared error attributed to that variable in a boosted regression tree model.

| Variable | Relative importance (reduction in squared error) |
| --- | --- |
| Average number of records per species | 78.5 |
| Species distribution modelling method | 14.8 |
| Spatial bias | 6.7 |

**Table 3.** Spatial sampling evenness of the spatial sampling template datasets measured using Simpson evenness. A value of one indicates perfectly even sampling (all grid squares containing the same number of records). Lower Simpson evenness values indicate more spatially uneven sampling.

| Spatial sampling template | Simpson evenness value |
| --- | --- |
| no bias | 1 |
| low bias (birds) | 0.762 |
| median bias (butterflies) | 0.126 |
| severe bias (moths) | 0.021 |

### 3.1 Number of species successfully modeled

The number of species for which models fitted successfully increased as sample size increased and spatial bias decreased (Fig. 4). For GLMs and inverse distance-weighted interpolation, model fitting was largely successful when datasets had more than 100 records per species, except when spatial bias was severe (Fig. 4). Boosted regression trees failed to fit models for some species even with relatively large amounts of data (e.g. an average 200 records per species), and models fit less frequently when data had median or severe spatial biases (Fig. 4). The effect of spatial bias on the number of species for which models fitted was small, but was slightly greater for boosted regression trees than for other SDM modelling methods (Fig. 4).

**Fig. 4.**
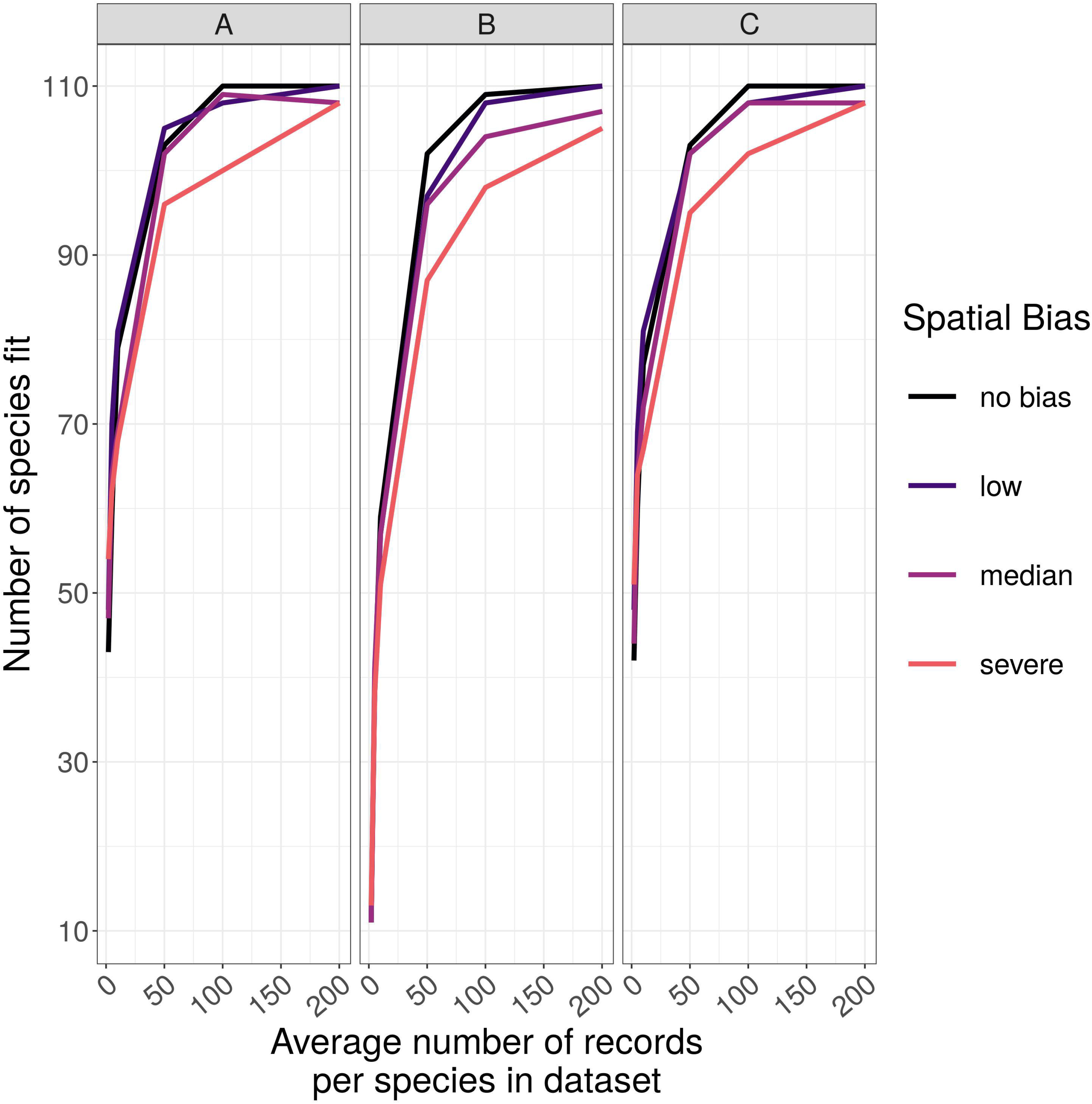
The number of virtual species successfully modeled. The number of virtual species (out of 110 total species chosen for modelling from the large community simulation) for which species distribution models fitted within the computation time and memory constraints we imposed, according to the spatial sampling bias and sample size of training data and the species distribution modelling method. Species distribution modelling methods were (A) generalized linear models, (B) boosted regression trees, and (C) inverse distance-weighted interpolation. Spatial biases were no bias (Simpson evenness = 1), low (e.g. birds, Simpson evenness = 0.76), median (e.g. butterflies, Simpson evenness = 0.13), and severe (e.g. moths, Simpson evenness = 0.02).

### 3.2 Predictive performance of SDMs

The amount of spatial bias in training data was less important than sample size and choice of SDM method in predicting the performance of SDMs (Table 2, Table S7, Table S8). AUC for predictive SDMs increased with the average number of records per species and with decreasing spatial bias in the training data when using all SDM methods (Fig. 5, Fig. 6). Root mean squared error (RMSE) was largely unaffected by spatial sampling bias (Fig. 7, Fig. S6, Table S8). Species prevalence (the number of grid squares occupied by a species) and the number of detections in the test dataset both had negligible effects on the average value of AUC, though they did affect the variability of AUC (Fig. S4, Fig. S5). Species prevalence strongly affected the expected value of RMSE, with RMSE increasing with species prevalence (Table S8, Fig. S4).

**Fig. 5.**
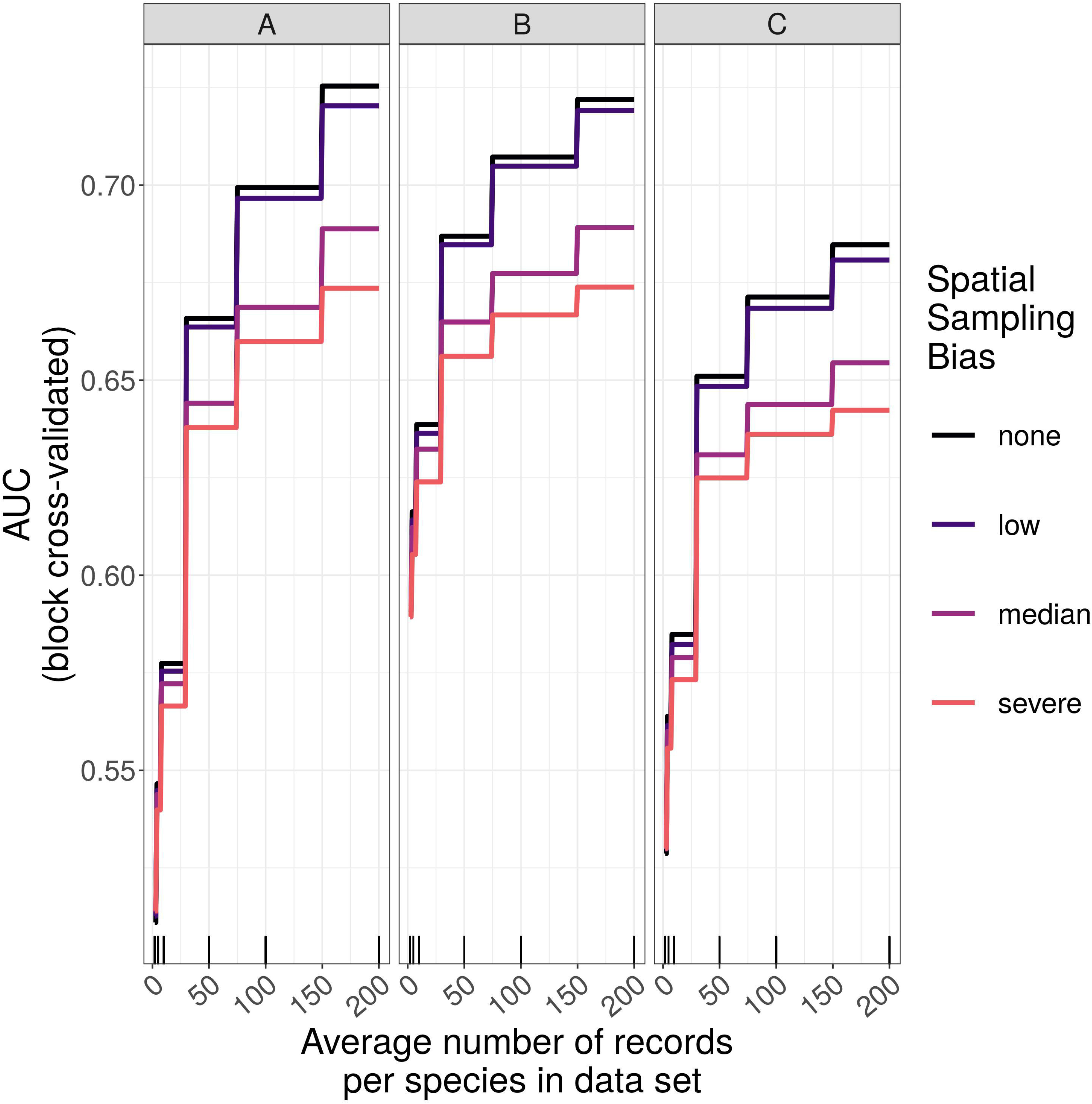
Expected prediction performance of species distribution models for 110 simulated species under a range of sample size and spatial sampling bias scenarios. Panels show the expected prediction performance of species distribution models constructed using (A) generalize linear models, (B) boosted regression trees, and (C) inverse distance-weighted interpolation. Lines show expected area under the receiver operating characteristic curve (AUC) given the sample size and spatial sampling bias of training data, and the species distribution modelling method. Rug plots indicate sample sizes (mean number of records per species) of the virtual biological records datasets used to train species distribution models.

**Fig. 6.**
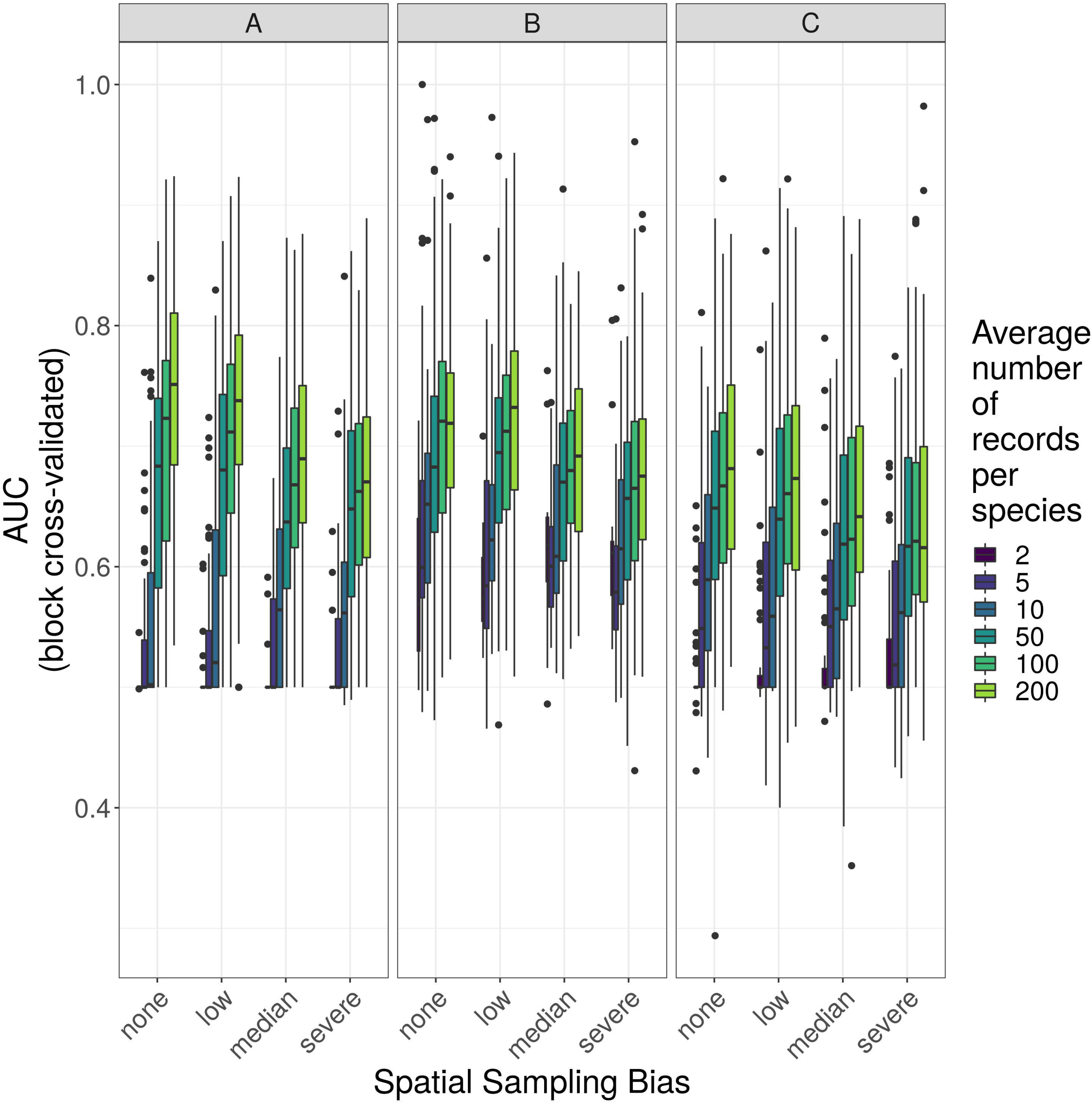
Observed prediction performance (AUC) of species distribution models for 110 virtual species under a range of sample size and spatial sampling bias scenarios. Panels show the observed area under the receiver operating characteristic curve (AUC) of species distribution models constructed using (A) generalized linear models, (B) boosted regression trees, and (C) inverse distance-weighted interpolation. Boxes contain the middle 50% of the observed AUC values. The horizontal line within each box indicates the median AUC value. Each box plot (box, whiskers, and outlying points) represents 110 observations (one for each virtual species) unless models failed to fit for some species (see Fig. 4). The width of boxes is proportional to the square root of the number of observations in that group.

**Fig. 7.**
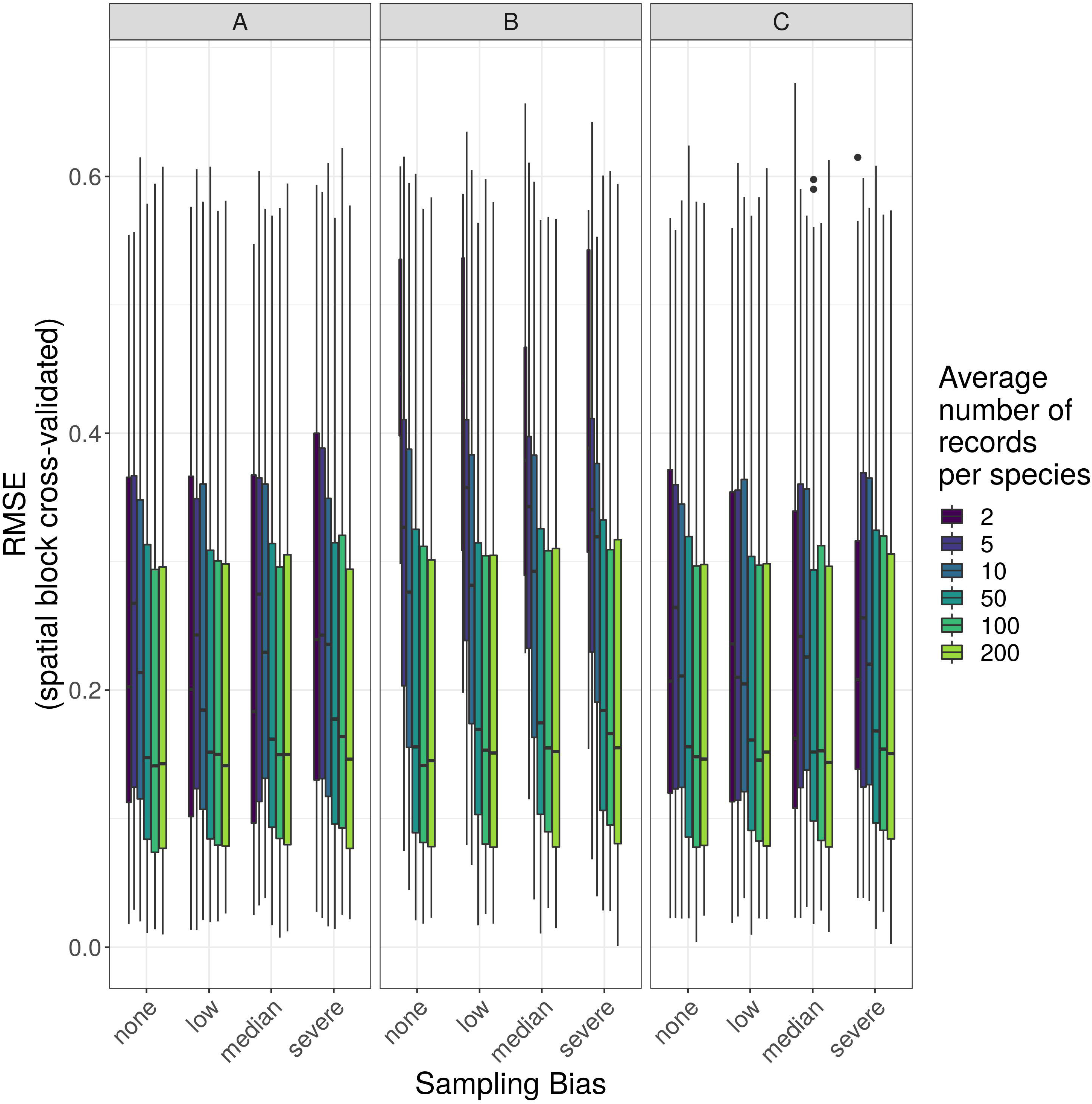
Observed prediction performance (RMSE) of species distribution models for 110 virtual species under a range of sample size and spatial sampling bias scenarios. Panels show the observed root mean squared error (RMSE) of species distribution models constructed using (A) generalized linear models, (B) boosted regression trees, and (C) inverse distance-weighted interpolation. Boxes contain the middle 50% of the observed RMSE values. The horizontal line within each box indicates the median RMSE value. Each box plot (box, whiskers, and outlying points) represents 110 observations (one for each virtual species) unless models failed to fit for some species (see Fig. 4). The width of boxes is proportional to the square root of the number of observations in that group.

#### 3.2.1 Effect of sample size

Sample size (average number of records per species) was the most important variable for predicting species distribution model prediction performance (Table 2). AUC improved with increasing average number of records per species for all SDM methods, and the improvement in AUC decelerated as the number of records per species increased (Fig. 5, Fig. 8).

**Fig. 8.**
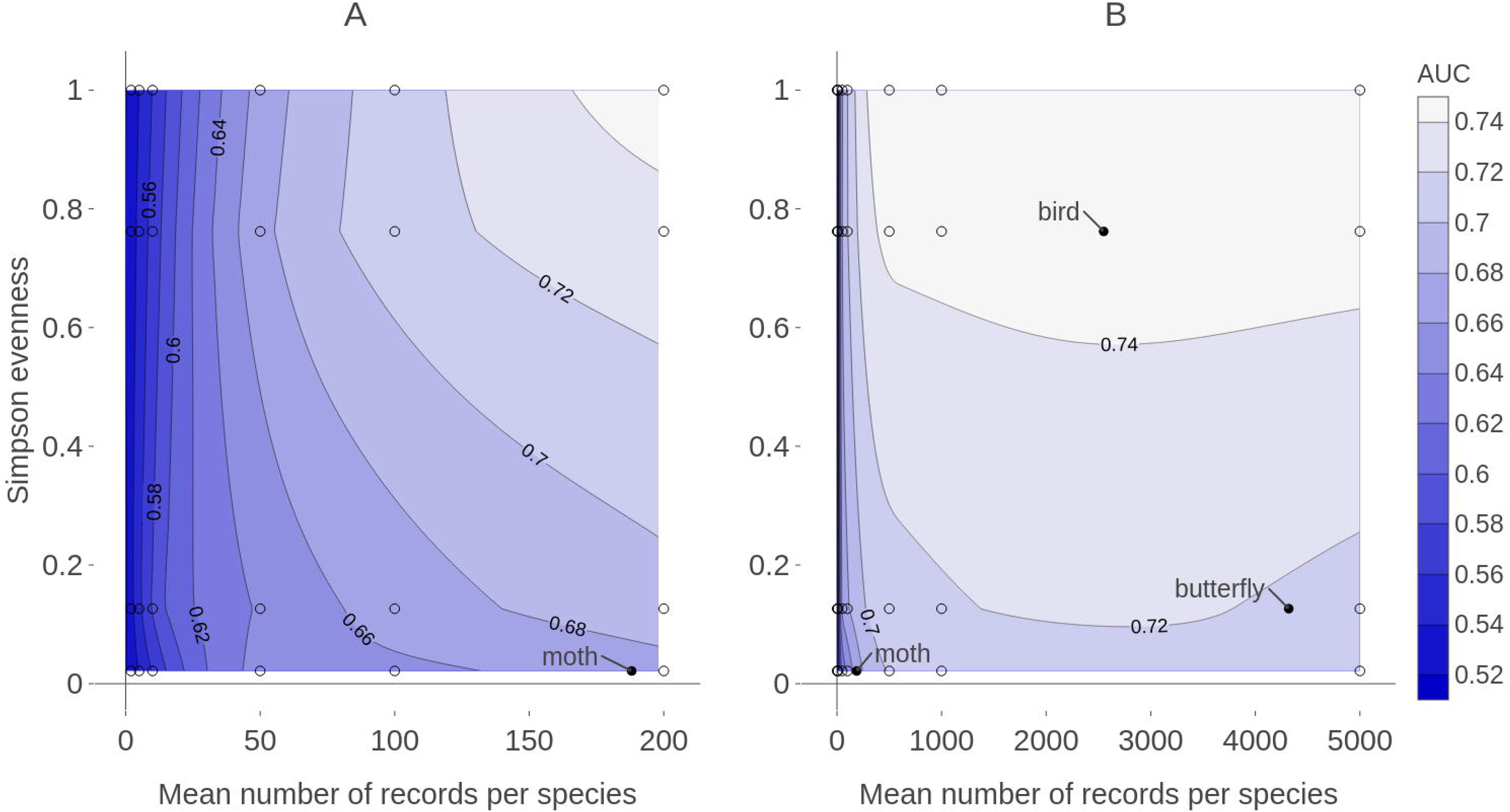
Contour plot of expected prediction performance of species distribution models as a function of the sample size and spatial sampling bias in virtual biological records datasets. Expected prediction performance (AUC, contours and shading) of generalized linear model (GLM) species distribution models from the (A) large- and (B) small-community simulations, according to the spatial sampling evenness and sample size of training data (note the different scales of the horizontal axes in A and B). Spatial sampling evenness was quantified using Simpson evenness. High values of Simpson evenness indicate minimal spatial bias. Open circles show the values of sample size and spatial sampling evenness for virtual biological records datasets used to train species distribution models. Filled black circles show sample size and spatial sampling evenness of Irish biological records datasets used as spatial sampling templates.

#### 3.2.2 Effect of spatial bias

Higher levels of spatial sampling bias generally reduced AUC, but the size of this effect was small for the low level of bias (Fig. 5). SDMs built with GLMs showed the biggest difference in prediction performance between models trained with unbiased data and models trained with data showing median spatial bias (reduction in expected AUC of 0.037 when using an average of 200 records per species, Fig. 5). Other SDM methods showed less difference in AUC between models trained with unbiased data and models trained with data containing median spatial bias (decrease in expected AUC of 0.033 for boosted regression trees and 0.030 for inverse distance-weighted interpolation when using an average of 200 records per species).

The AUC for inverse distance-weighted interpolation models trained with unbiased data was generally higher than the AUC for GLMs and boosted regression trees trained with severely biased data, but lower than the AUC for GLMs and boosted regression trees trained with data with median spatial bias for any given sample size (Fig. 5, Fig. 6).

## 4 DISCUSSION

Both sample size (the average number of observations per species) and choice of modelling method were more important than the spatial bias of training data for determining model prediction performance. This is in line with the results of Thibaud et al. (2014). However, Thibaud et al. (2014) simulated spatial sampling bias by defining sampling probability as a linear function of distance from the nearest road. In contrast, our study used observed spatial sampling patterns from real biological records datasets. Our results therefore provide a more direct confirmation that spatial biases of the type and intensity found in real datasets are not as important as other factors in determining SDM prediction performance.

While spatial bias was not the most important factor determining SDM prediction performance, spatial sampling bias did affect model prediction performance when spatial bias was relatively strong. The limited effect of spatial bias on SDMs that we observed is similar to other findings that have shown spatial sampling bias to have a small effect on model performance (Thibaud et al., 2014; Warton et al., 2013) or to affect only some SDM methods (Barbet-Massin et al., 2012). Given Fan et al.’s (2005) conclusion that most types of predictive models can be either sensitive or insensitive to sample selection bias in training data, depending on the specific datasets, it seems unlikely that a broad conclusion about the effect of spatial sampling bias on species distribution models in all cases is possible. It therefore remains important to test the effect of spatial bias on SDMs using data that match as closely as possible the data used for different SDM applications. Our study used spatial biases and the spatially explicit environmental data representative of data likely to be used in SDMs using biological records in Ireland. Our conclusions therefore apply most directly to applications of SDMs using Irish biological records, and may not be generalizable to other geographic locations, or for species within Ireland that do not respond to the environmental predictor variables used in this study. However, our results strengthen a growing body of literature that suggests that spatial sampling bias is rarely the most important issue in determining SDM prediction performance. In particular, the choice of modelling method may often have more impact on SDM prediction performance than a variety of other factors (Barbet-Massin et al., 2012; Fernandes, Scherrer, & Guisan, 2018).

Training data with low spatial sampling bias produced species distribution models that performed nearly as well as models trained with unbiased data. Prediction performance was poor when models were trained with small sample sizes, regardless of the spatial bias in training data. Similarly, model performance increased quickly with sample size when sample size was small, even when the data had severe spatial bias. This suggests that, for taxonomic groups with relatively few records per species, the usefulness of the data for predictive SDMs can be improved by increasing sample size, even if additional data collection is spatially biased. In contrast, for taxonomic groups for which biological records datasets already have a high average number of records per species (e.g. birds and butterflies which both have an average of over 2000 records per species in Ireland) further improvements in SDM prediction performance will likely require increasing the spatial evenness of data (Fig. 8).

The objective of our SDMs was to fill in gaps in species distribution knowledge within the spatial and environmental conditions of the island of Ireland, an area of about 84,000 km^2^. Our results may not generalize to larger spatial scales or to cases in which the goal of SDMs is uncovering species’ entire fundamental environmental niche or determining the environmental factors most strongly influencing distributions. The spatial scope of our SDMs is sensible both from an ecological and applied standpoint, because the island of Ireland is a geographically delimited ecological unit, and because decision making about species conservation and management often happens within political units (e.g. nations, states, or counties) that cover only a portion of species’ spatial and environmental distributions. Our results suggest that, when the goal of predictive SDMs is to fill in data gaps within a scale of tens of thousands of square kilometers (e.g. a national scale in the case of Ireland), spatial sampling bias was less important in determining model performance than the total amount of data and the SDM modelling method.

GLMs had the best prediction performance of the four SDM methods we tested, even though they were more affected by spatial bias than were other methods. The high performance of GLMs relative to other modelling methods in this study agrees with the simulation results of Thibaud et al. (2014) and Fernandes et al. (2018). However, as in both those studies, we generated virtual species distributions according to a linear model, so it is possible that the good performance of GLMs is due to the model having the same functional form as the “true” species responses. In real applications, it is unlikely that the functional form of the model will exactly match the form of the true species responses. Indeed, the species distribution modelling literature has many examples of different modelling methods performing best in different studies, suggesting that no modelling method consistently outperforms others (Bahn & McGill, 2007; Breiner, Nobis, Bergamini, & Guisan, 2018; Cutler et al., 2007; Elith et al., 2006; Elith & Graham 2009).

Boosted regression trees’ prediction performance was slightly less affected by spatial bias than GLMs’, and prediction performance of both methods was similar when trained with large, spatially biased datasets. But boosted regression trees failed to fit models more often than did GLMs, especially when sample sizes were smaller, which may make them inferior to other modelling methods for small datasets, at least within the computational resource limits we faced. We cannot rule out the possibility that the performance of boosted regression trees would improve if they were trained with a smaller learning rate and permitted to grow more than 30,000 trees. However, most users of SDMs will face some computational resource limitations. We permitted boosted regression trees to grow up to 30,000 trees, which is well above the rule-of-thumb guidelines given by Elith, Leathwick, and Hastie (2008).

In this study, we introduced spatial bias specifically into the training data and tested model performance using spatially even evaluation data. However, spatial bias can also occur in evaluation data and may affect the reliability of model evaluations (Fink et al., 2010). When using real biological records datasets, it is likely that both model training and evaluation will use spatially biased data, making it difficult to dis-entangle whether observed effects of spatially biased data on prediction performance are due to the influence of biased data in the model training step or in the model evaluation step. We evaluated models on spatially even data (which is easy using simulated data but would be more difficult or impossible when using real data), so the observed effects of spatially biased data on prediction performance in our study can be attributed to the effect of biased data on model training. All of the SDM methods we used involve some kind of model evaluation as part of the model training process, either inherent in the model fitting or introduced by our implementation. For example, with our GLMs we introduced a model evaluation step when we chose the combination of predictor variables that gave the model with the lowest AIC on training data. The final GLM models were therefore based on variables that had been selected by evaluation on spatially biased data. For both GLMs and inverse distance-weighted interpolation, it is possible that using unbiased data in the evaluations during model selection would have led to different final models. Therefore, the observed effect of the spatial bias in this study could be due to how biased data affects the actual fitting of each individual model, or to how the biased data affects the evaluation step used to select which fitted model to use for predictions. Tree-based methods, including boosted regression trees, select which values of predictor variables to split at and/or which predictor variables to use at each node based on how much those splits improve some measure of performance on the training data (Elith et al., 2008; Hastie et al., 2009). Thus, evaluation on potentially spatially biased training data is inherent in fitting tree models.

Fink et al. (2010) provided a method for correcting spatial bias in evaluation data to reduce the effect of spatial bias on model evaluation, but they did not explicitly address spatially biased data in model training. Our results showed that spatially biased data can impact model training (at least when the spatial bias is relatively strong). Investigating the effect of spatially biased data on the evaluation that takes place as part of model training (e.g. during variable selection or parameter tuning) may be a worthwhile path for future research. It may be possible to use a method like that proposed by Fink et al. (2010) to correct spatial bias during the evaluation that takes place within the model training process. This may reduce the effect of spatially biased training data on model performance that we observed.

Our use of Simpson evenness to measure spatial sampling evenness allows the spatial sampling biases tested in this study to be compared to spatial sampling patterns in existing datasets. Because we calculated spatial sampling evenness using the number of records in each grid square relative to the entire study extent, our measures of spatial sampling evenness confound species richness and sampling effort. Using the number of checklists (or sampling events) rather than the number of records would alleviate this problem. However, records in our datasets were aggregated over long time periods so that the records appear to have the same date, location, and observer, even when records arose from different sampling events. For example, records from vascular plant and bird atlases have been incorporated into the NBDC database with all the atlas records from a grid square being assigned the same date (the publication date of the atlas), even though records were collected over multiple years. Many of these atlas grid square “checklists” are hundreds (or thousands!) of records long, with repeat observations of common species. The total number of records therefore better represents the many years and many unique days of sampling in heavily sampled grid squares for NBDC datasets, despite the fact that spatially uneven species richness will cause the number of records to be higher in some grid squares than others, even when sampling effort is equal.

## 5 CONCLUSION

We found that spatial sampling bias in training data affected species distribution model prediction performance when the spatial bias was relatively strong, but that sample size and the choice of modelling method were more important than spatial bias in determining model prediction performance. This study adds to a body of literature suggesting that prediction performance of species distribution models is less affected by spatial sampling bias in training data than by other factors including modelling method and sample size.

## 6 ACKNOWLEDGMENTS

We thank Tomás Murray and the Irish National Biodiversity Data Centre (NBDC) for providing and answering questions about the biological records data, and we thank the many citizen scientists who collected and contributed their data to the NBDC. This publication emanated from research supported in part by a grant from Science Foundation Ireland under grant number 15/IA/2881. This work used the ResearchIT Sonic cluster funded by UCD IT Services and the Research Office. This research used CORINE data made available with funding by the European Union.

