## Supplementary figures and images for "Data quantity is more important than its spatial bias for predictive species distribution modelling"

### Supplemental Figure S1

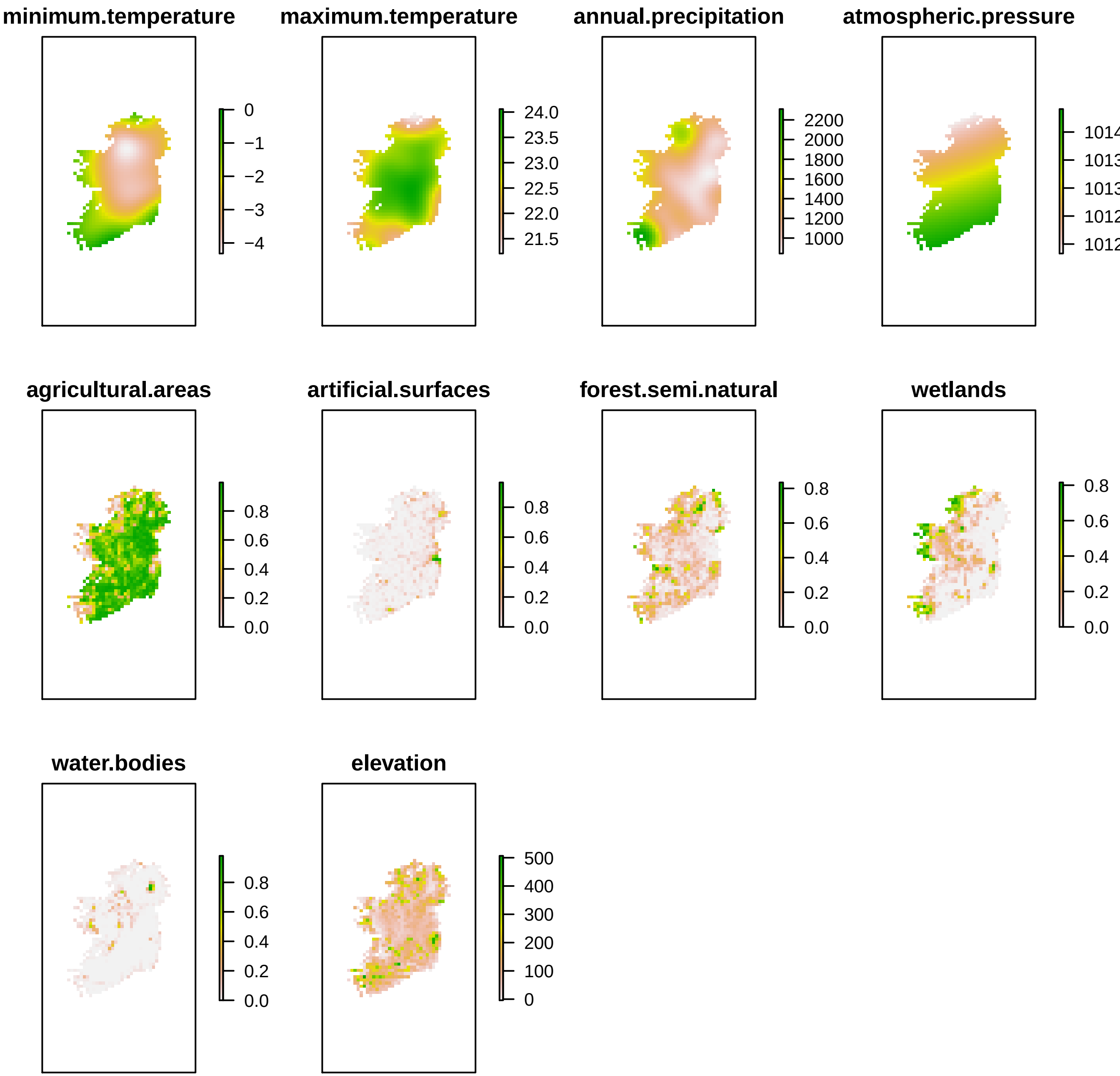

### Supplemental Figure S4

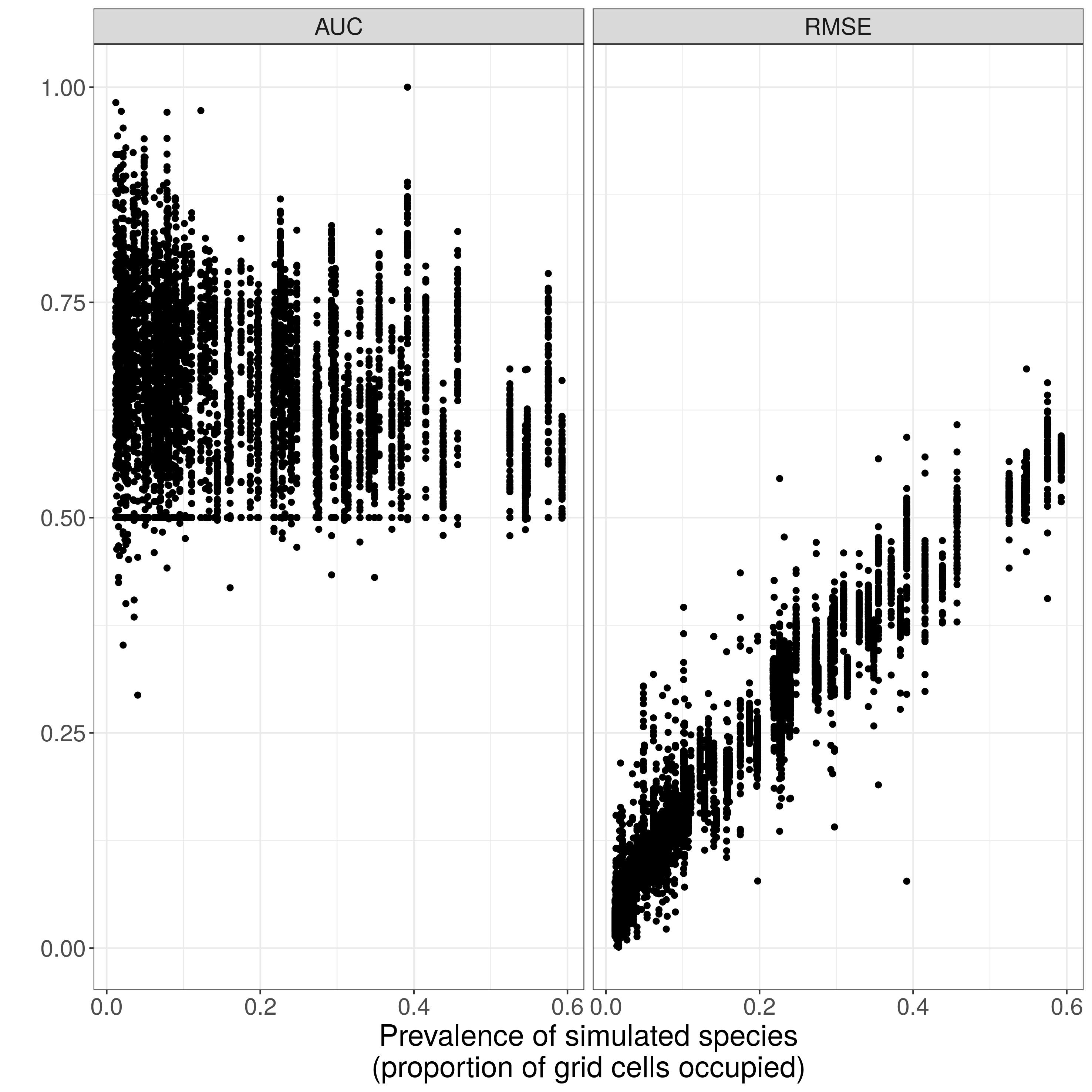

### Supplemental Figure S5

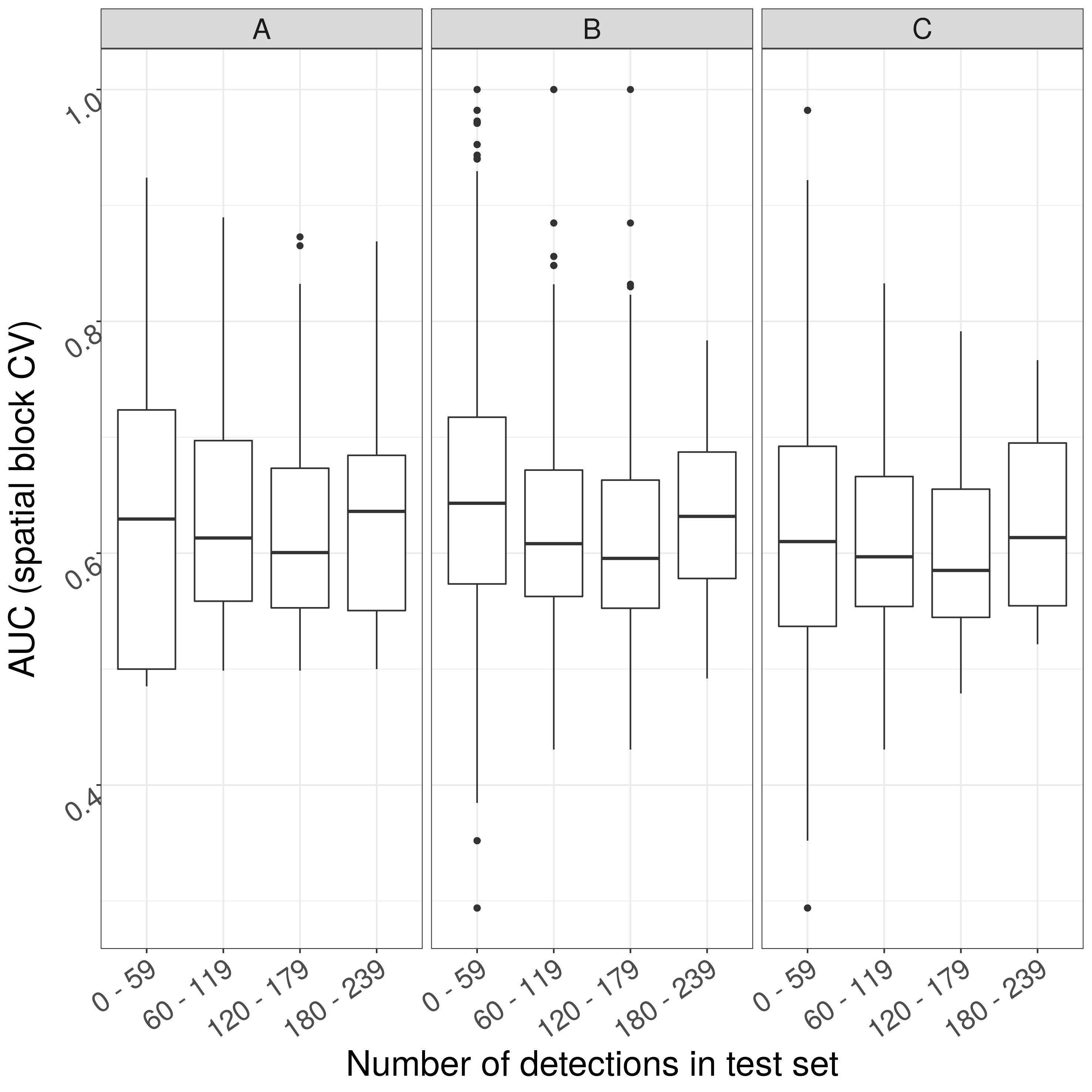

### Supplemental Figure S6

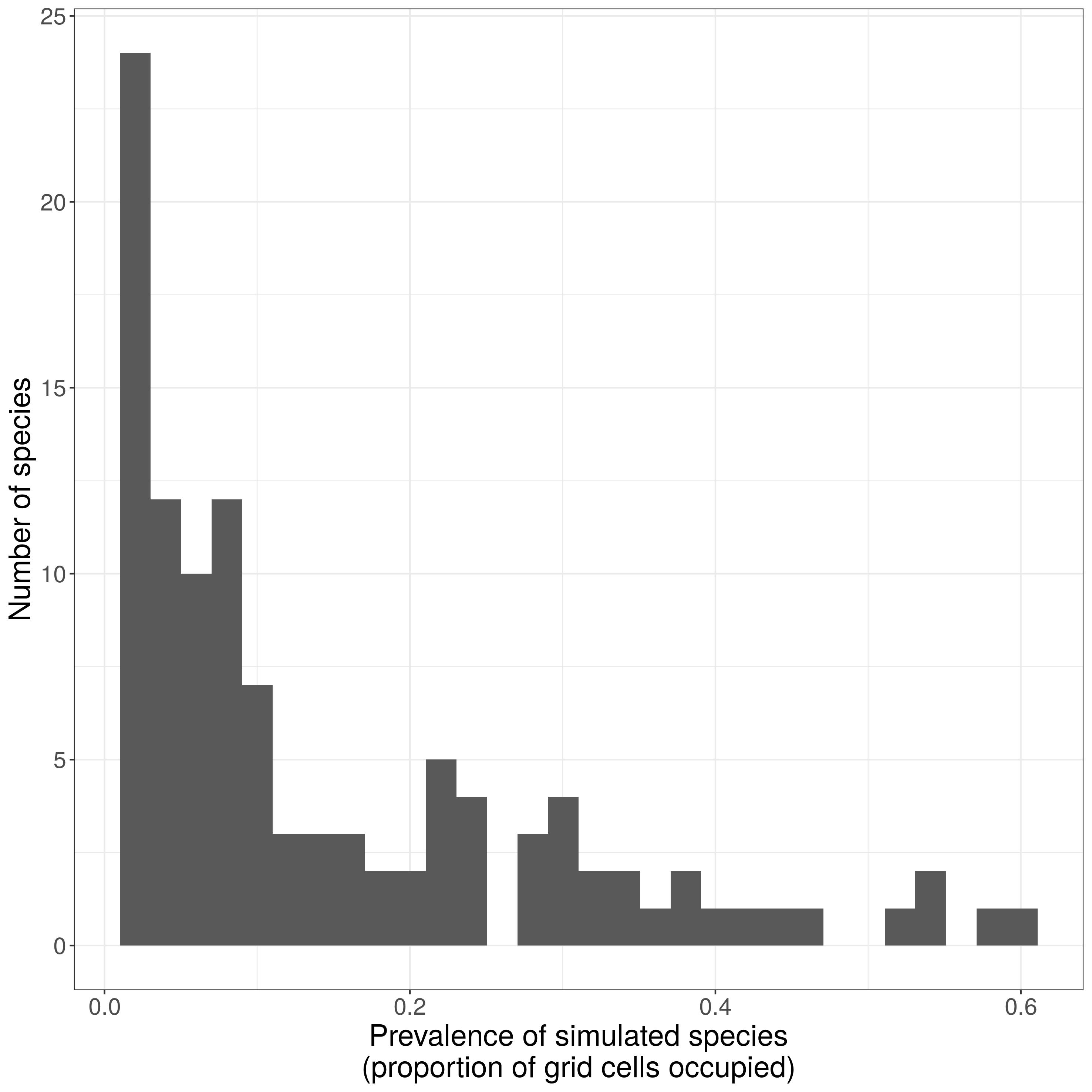

### Supplemental Figure S9

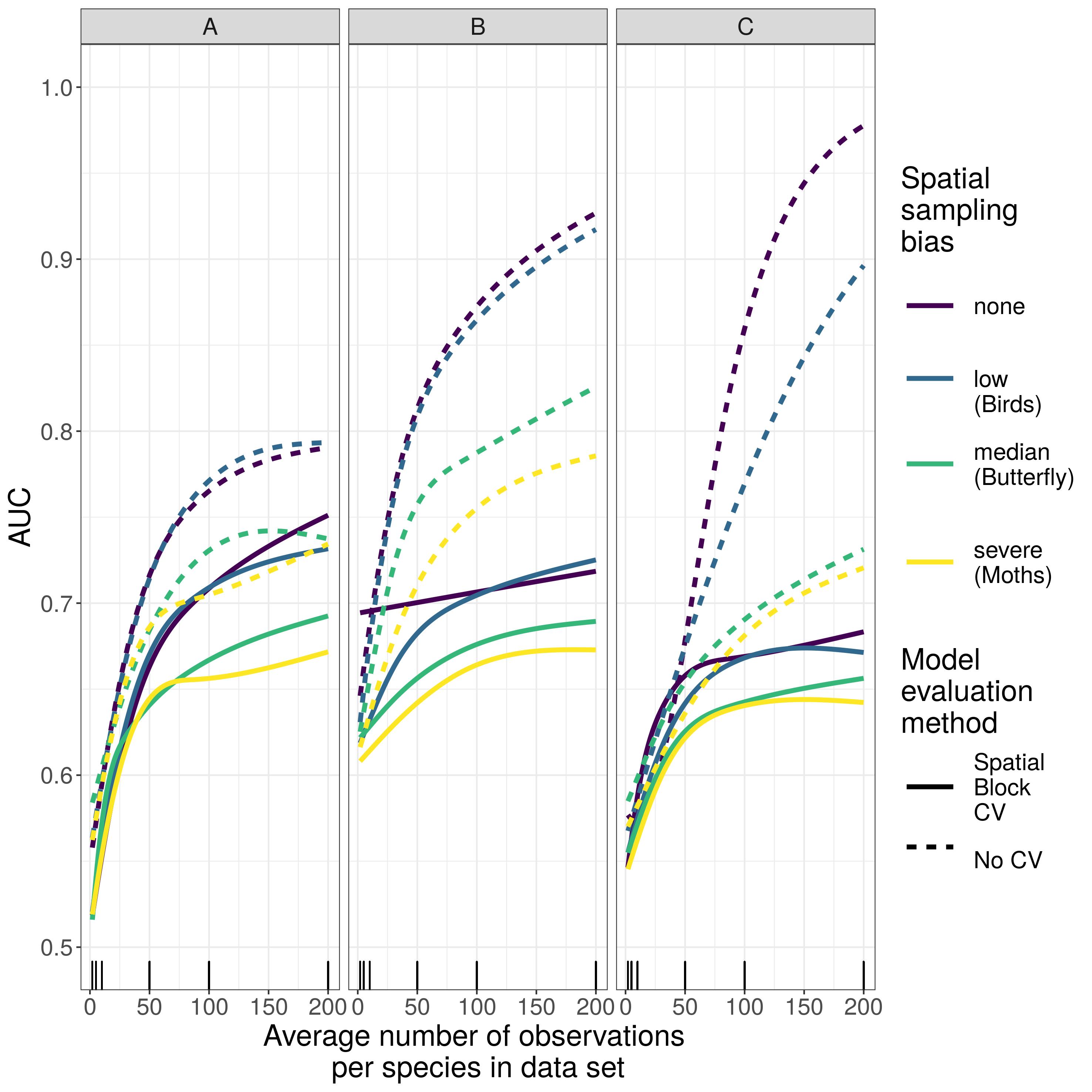

### Supplemental Figure S10

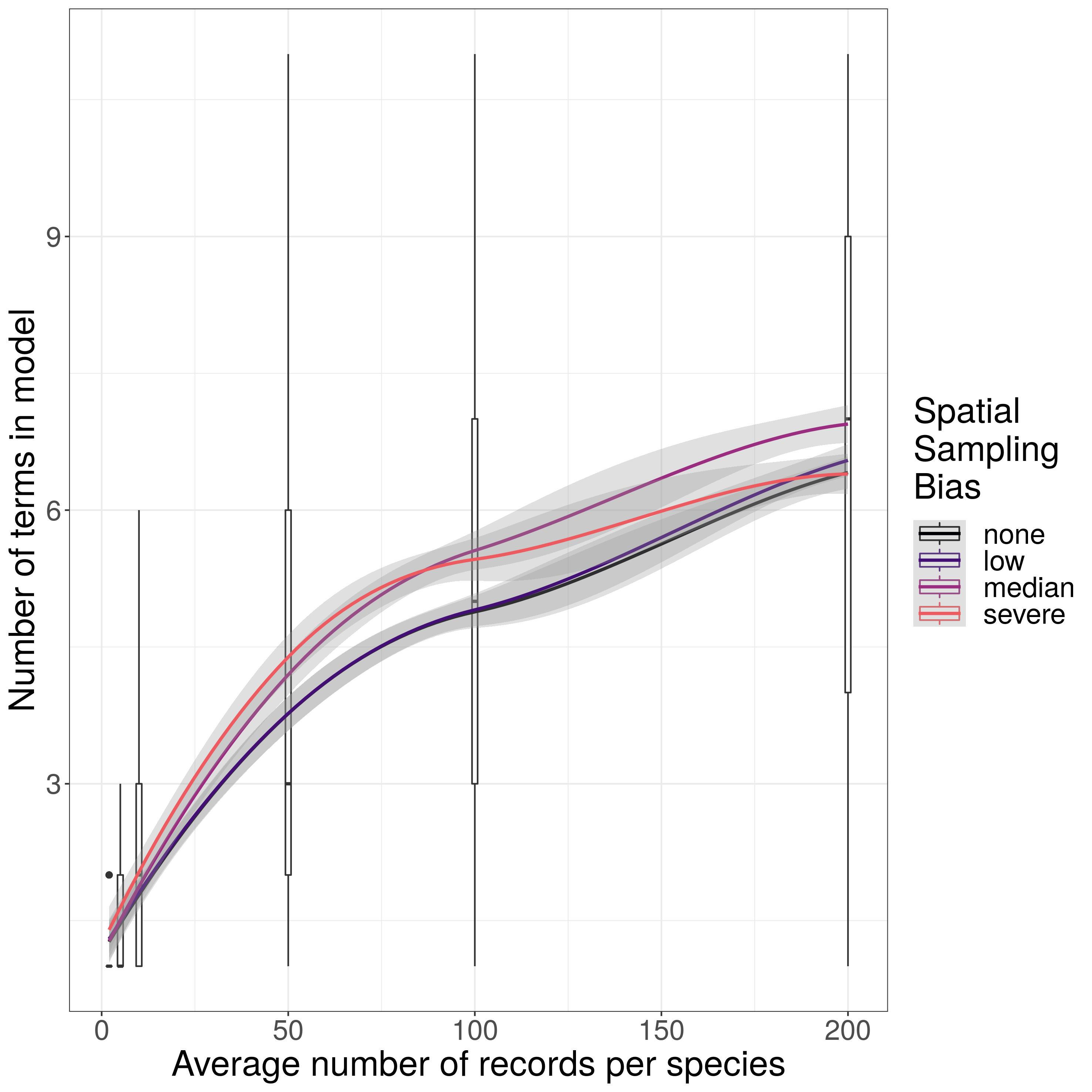

### Supplemental Figure S11

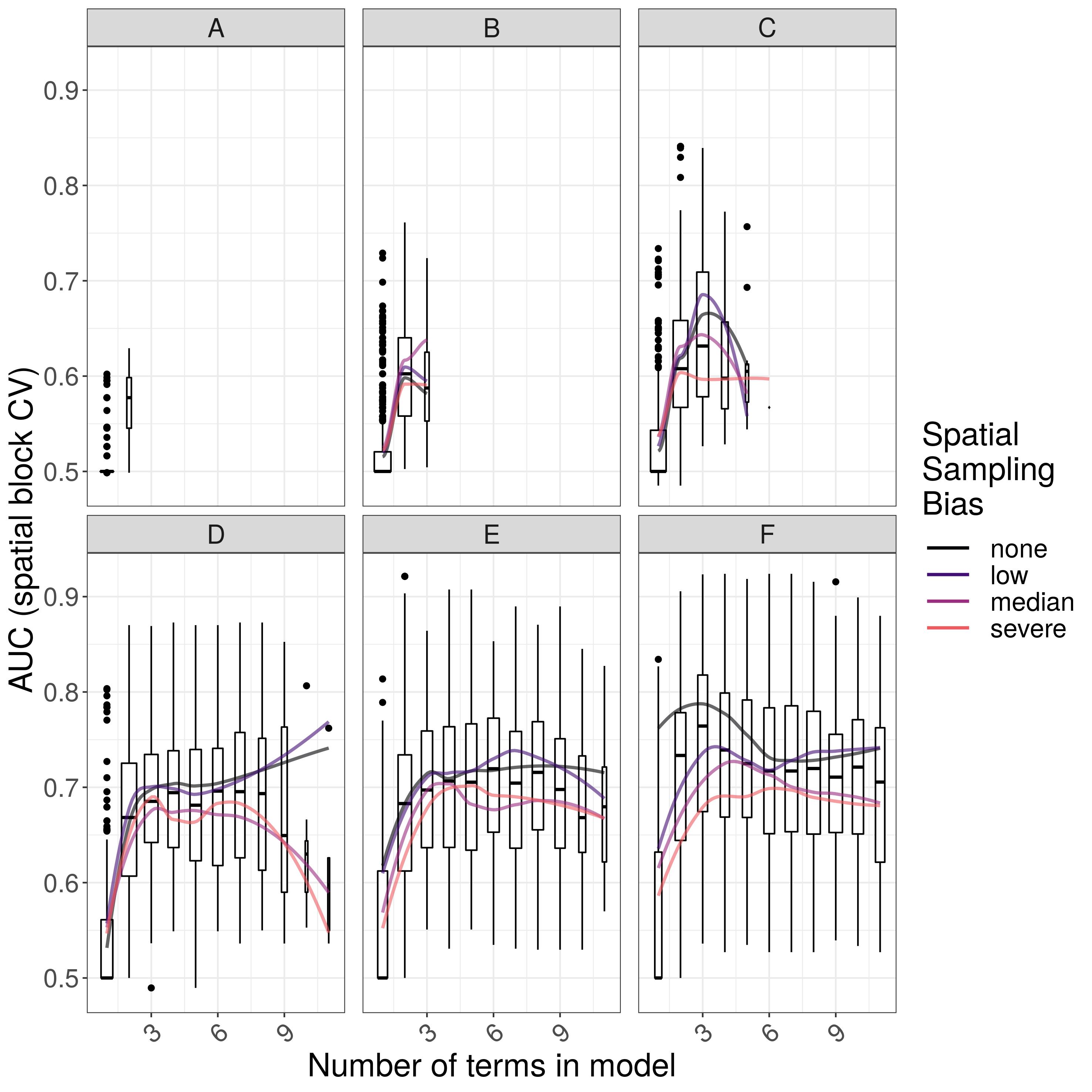

### Supplemental Figure S12

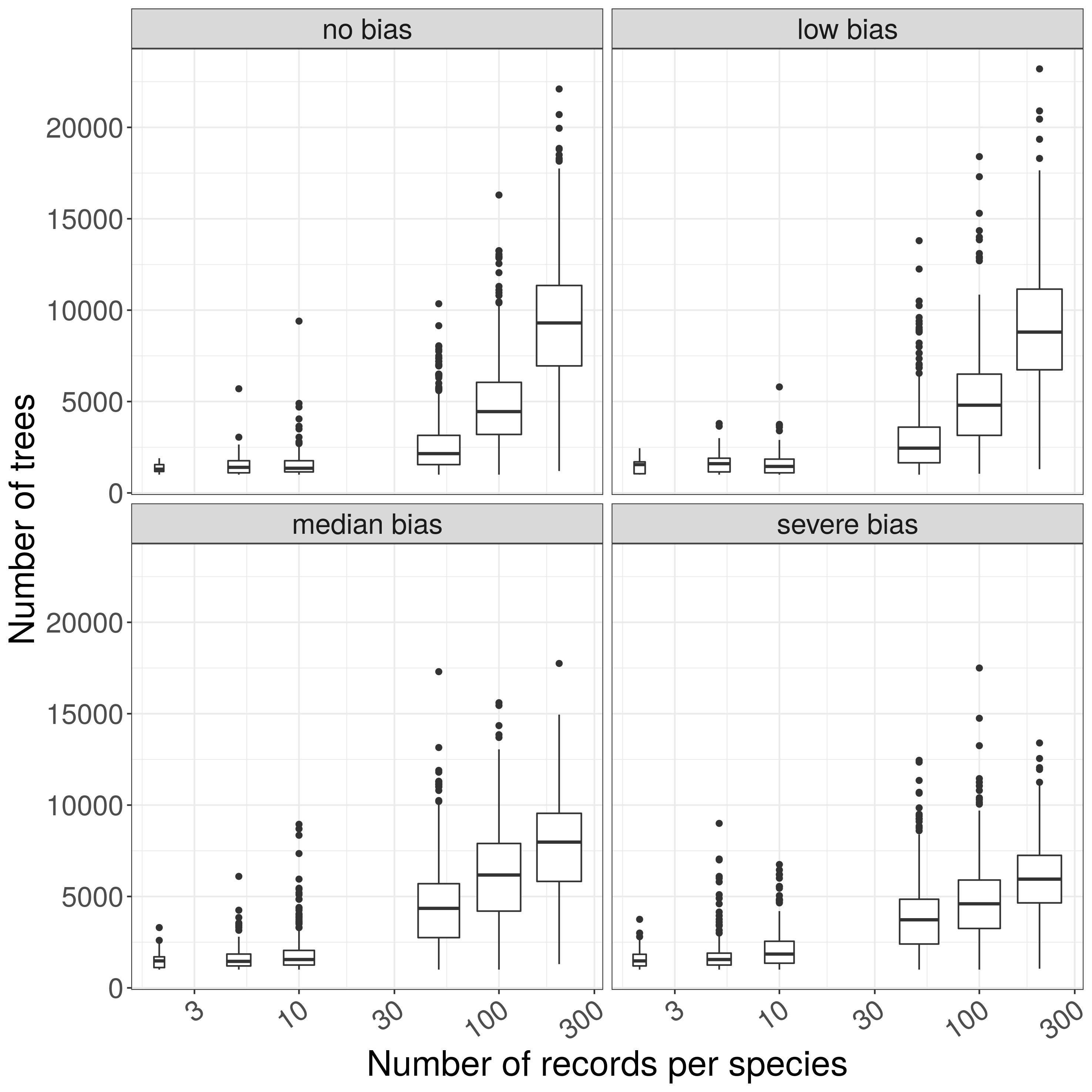

### Supplemental Figure S13

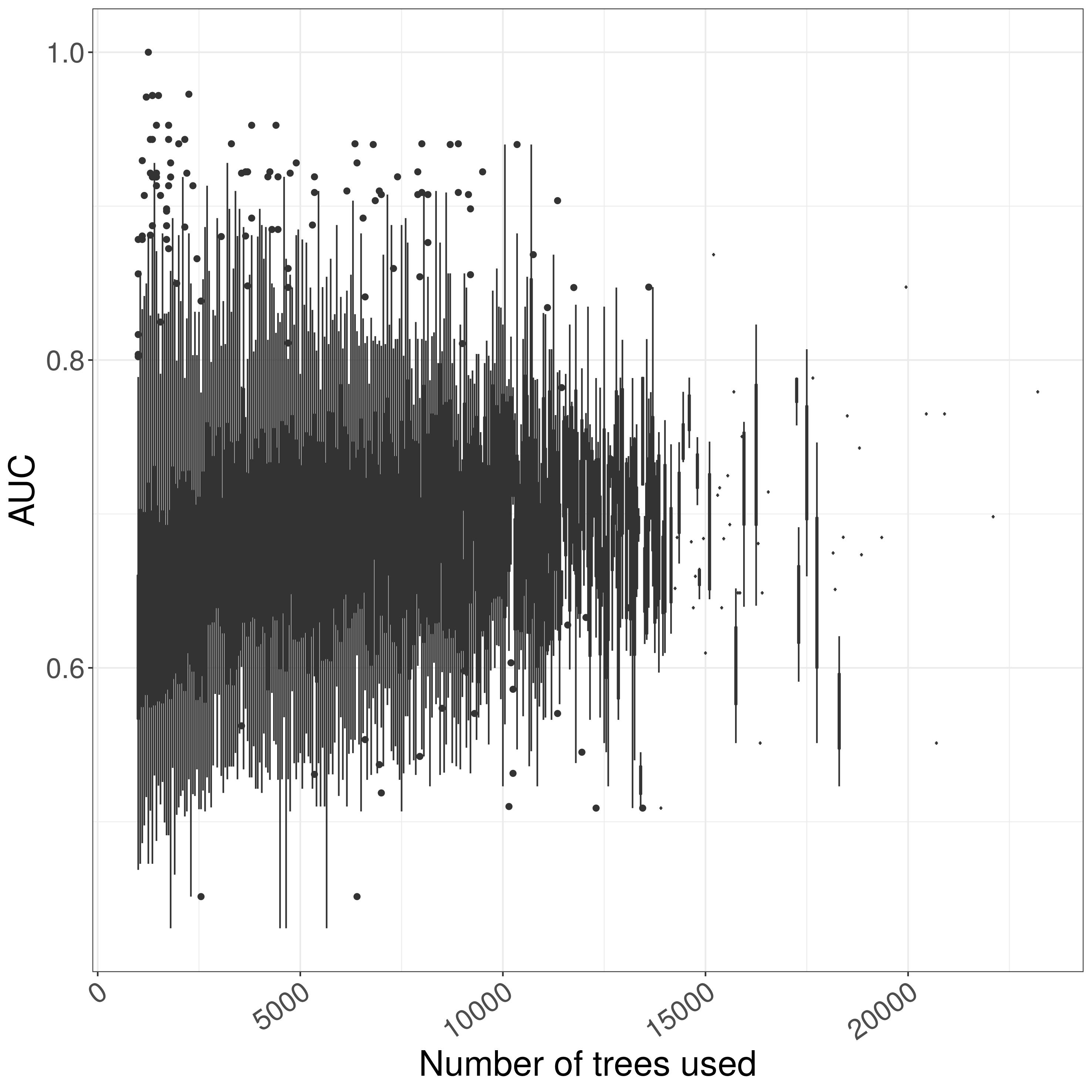

### Supplemental Figure S14

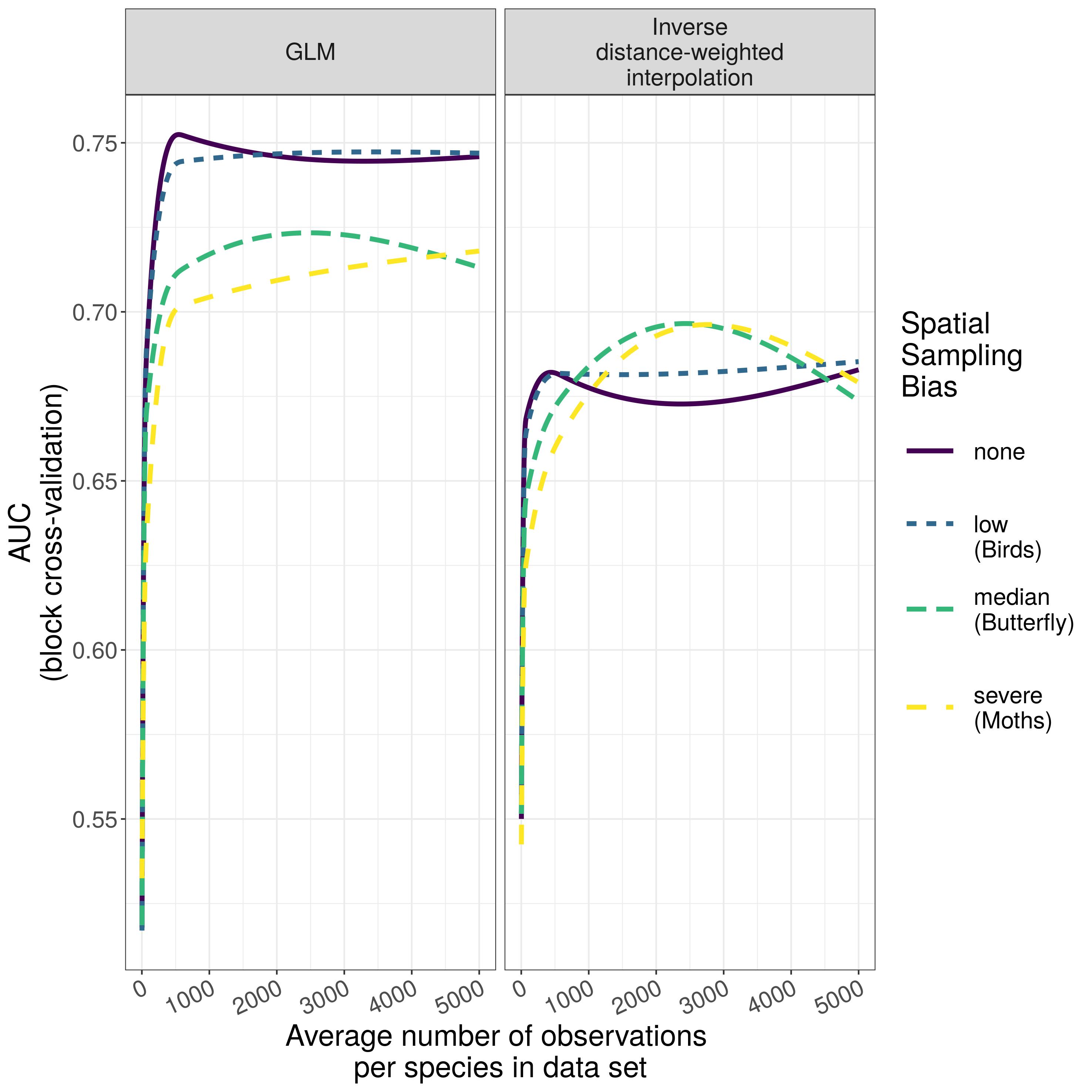
