## Supplemental Text S2 for "Data quantity is more important than its spatial bias for predictive species distribution modelling"

### S2: Supplementary Methods and Results

Willson Gaul[1*], Dinara Sadykova[2], Hannah J. White[1], Lupe León-Sánchez[2], Paul Caplat[2], Mark C. Emmerson[2], Jon M. Yearsley[1]

1. School of Biology & Environmental Sciences, University College Dublin, Dublin, Ireland

2. School of Biological Sciences, Queen's University Belfast, Belfast, UK

[remainder of this page intentionally blank]

### Supplementary methods

#### Environmental Predictor Variables

We used average minimum annual temperature, average maximum annual temperature, average annual precipitation, and average daily sea level atmospheric pressure calculated over 12 years (1995 to 2016) from the E-OBS European Climate Assessment and Dataset EU project (Haylock et al., 2008; van den Besselaar, Haylock, van der Schrier, & Klein Tank, 2011; http://www.ecad.eu/download/ensembles/downloadchunks.php). For average minimum and maximum temperatures, we calculated the mean across all 12 years of the 2% and of the 98% quantiles of daily mean temperatures. For average annual precipitation, we summed daily precipitation within each year and calculated the mean annual precipitation over all years (excluding years 2010 through 2012 because of missing daily precipitation values in those years). For average daily sea level pressure we took the mean of daily sea level pressure over all 12 years. We calculated the value of each climate variable at the E-OBS grid points and then interpolated to Irish 10 km grid cells using ordinary kriging.

We calculated the proportion of each grid cell covered by each of the “agricultural areas”, “artificial surfaces”, “forest and semi-natural areas”, “water bodies”, and “wetlands” Label 1 categories from the CORINE Land Cover database (CORINE, 2012). We calculated the average elevation within each grid cell by interpolating using ordinary kriging from the ETOPO1 Global Relief Model (Amante & Eakins, 2009; https://www.ngdc.noaa.gov/mgg/global/relief/ETOPO1/data/ice_surface/grid_registered/netcdf/ [accessed 8 May 2019]).

Spatial clustering of predictor variable values was measured using Moran’s I calculated with the ‘Moran’ function in the ‘raster′ R package (Hijmans, 2018).

**Simulating species distributions**

Coefficients specifying the virtual species' responses were chosen such that the theoretical prevalence of the species (the sum of the probabilities of presence in each grid square divided by the number of grid squares) was greater than 0.01 so that virtual species were common enough to be observed and modeled. Coefficients for the squared terms were randomly drawn from a uniform distribution between zero (which creates a straight-line response) and 1.3 (chosen because higher values produce response curves with narrow "humps", representing species with highly specialized environmental niches such that there would be very few occurrences within Ireland). The maximum coefficient value of 1.3 for the squared terms was chosen after exploring multiple values, with the goal of finding a value that regularly produced virtual species with theoretical prevalences across the entire study extent greater than 0.01 (corresponding to the species being present in about 8 grid cells in Ireland). Coefficients for $1^{st}$ order terms were randomly drawn from a uniform distribution within minimum and maximum values chosen to ensure that the response to each predictor variable had an optimum within the range of values of the predictor variable within Ireland.

**Simulating sampling with spatial bias**

The reason for varying the probability of sampling a species according to species prevalence (Section 2.4.3 of main text) was to simulate the real-world scenario in which species that are present in many locations also likely have higher abundances (Gaston et al., 2000) and are therefore more likely than rare species to be recorded in any single sampling event. We defined the probability of observing a species as the twentieth root of that species's prevalence in the entire study extent. The twentieth root was chosen based on exploratory trials in which we generated checklists with species sampling probability weights defined by different transformations of prevalence (e.g. raw prevalence, square root of prevalence, or logarithm of prevalence) and looked at histograms of the number of observations per species and scatter plots of the number of observations of species by the prevalence of species. For many transformations of prevalence, including the logarithm of prevalence and the square root of prevalence, weighting sampling probability by the transformation generated sampled species lists that seemed badly unrealistic (e.g. weighting by raw prevalence produced checklists with almost only common species, and weighting by the natural logarithm of prevalence produced checklists of mostly rare species). Weighting by the twentieth root produced sampled species lists that seemed plausibly realistic in terms of the relative numbers of rare and common species sampled. Determining the probability of observing a species based on the species's prevalence in the overall study extent meant that the probability of observing a species when it was present was the same across the entire study extent.

Because we sampled occurrence records with replacement from the list of present species, it was possible for a species to appear on a sampling event checklist more than once. This matched the nature of many NBDC datasets in which some sampling event checklists were aggregations of records over long periods of time (e.g. all records from a location in a single year were aggregated and reported with an identical location and date). In those cases, a sampling event checklist may contain hundreds of records with many repeat observations of some species.

**Species distribution modeling**

Models were fitted with both five-fold block cross-validation and with no cross-validation (evaluating on the training data). Using block cross-validation is best practice, so only those results are presented in the main text. We included fitting with no cross-validation to confirm that prediction performance measures appear overly optimistic when evaluation is done without cross validation (as has been reported in the literature) (Roberts et al., 2017).

Spatial block cross-validation (Roberts et al., 2017) partitioned the study extent into spatial blocks of 100 km x 100 km and then allocated each 100 km x 100 km block to one of five cross-validation partitions. The spatial position of the 100 km x 100 km blocks was determined randomly (by randomly setting an origin point for the grid). The exact number of 100 km x 100 km blocks required to cover the island of Ireland depended on the randomly-determined location of the grid cells. A 100 km x 100 km block could be (and often was) positioned partially over ocean. Therefore, not every 100 km x 100 km block contained the same number of terrestrial grid cells, and consequently not every block cross-validation fold contained the same number of terrestrial grid cells. Prediction performance (AUC and RMSE) of models was evaluated against true simulated species distributions at locations not included in the training set for each of the cross-validation folds, and AUC and RMSE values for all five cross-validation folds were averaged to produce the final values of AUC and RMSE describing the prediction performance of each model.

We provided five predictor variables to SDMs to model each species. The five predictor variables were chosen randomly from the 10 possible predictors (Table 1) in order to simulate a real-world situation in which the factors that influence species distributions are not entirely known, and variables used for modelling likely include a mix of important and unimportant variables. For GLMs, not all five predictor variables were necessarily used in the final model because of our model selection process (see below). All models used equal weights for presences and absences.

For the small community simulations, we fit models to 110 virtual species by creating three small communities, each with 40 virtual species (the number of recorded butterfly species in Ireland) and modelling all virtual species in each community except for the last community (from which we only modeled 30 virtual species).

*GLM*

We used a logistic regression ('glm' function in R) with a binomial error distribution and logit link to model the probability of a species being recorded during a sampling event. Quadratic terms for each of the five environmental predictor variables were fitted, but we did not fit interactions between variables. Within each of the five block cross validation (CV) partitions, we tested all possible models that contained few enough terms that there were at least 10 detections (or non-detections, whichever was smaller) per non-intercept term in the model. We chose as the final model the combination of predictor variables that gave the model with the lowest AIC based on the training data in that partition. Thus, the minimum model size was an intercept-only model, and the most complex model included an intercept plus 10 additional terms (1$^{st}$ and 2$^{nd}$ degree terms for each of the five predictor variables chosen to model that species). First degree terms for a variable were always included if a second degree term was selected for that variable. Because the goal was to produce predictive models for a large number of species, we did not assess model assumptions for each individual species model.

*Boosted regression trees*

We trained boosted regression trees using the function 'gbm.step' in the 'dismo' R package (Greenwell, Boehmke, & Cunningham, 2018; Hijmans, Phillips, Leathwick, & Elith, 2017). We tested models with tree complexities of two and five. Smaller learning rates are generally preferred because they result in better predictive performance but higher computation and memory requirements (Elith, Leathwick, and Hastie 2008). Therefore, for each tree complexity (two and five), we first tried to train each model with a learning rate of 0.001. If the model used fewer than 1000 trees, we shrank the learning rate by 50% in order to try to get models that used of 1000 trees (as recommended by Elith, Leathcwick, and Hastie 2008). If no model could be fitted with more than 1000 trees and a learning rate of higher than 0.00001, we abandoned model fitting for that training dataset. We used gbm.step to determine the optimal number of trees for each model, based on monitoring the change in 10-fold cross-validated error rate as trees were added to the model (Hijmans, Phillips, Leathwick, & Elith, 2017). We started models with 50 trees and added trees in increments of 50 (step.size = 50). If the model using the initial learning rate of 0.001 did not reach minimum error with fewer than 30,000 trees, we increased the learning rate incrementally by 0.002 until either the model fit successfully with fewer than 30,000 trees or the learning rate got larger than 0.1. If no model could be fit with fewer than 30,000 trees and a learning rate smaller than 0.1, we abandoned model fitting for that training dataset. Finally, we compared the models fit with tree complexities of two and five, and the optimum learning rate and number of trees selected for each of those models. Of the two models with different tree complexities, we chose as the final model the one that had the lower cross-validation predictive deviance. We then generated SDM predictions using this final model.

*Inverse distance-weighted interpolation*

Within each grid cell, we calculated the proportion of sampling events on which the focal species was recorded. For each CV partition, we used the 'gstat' function in R (Gräler et al., 2016; Pebesma, 2004) to predict the probability of recording the species during a sampling event in locations in the test partition by taking an inverse distance weighted average of proportions of training partition sampling events on which the species was recorded. The 'gstat' arguments specifying the optimal power parameter and number of points to use were chosen in an automated way by testing all combinations of powers in increments of 0.5 between 0 and 10 and number of points in increments of two between one and the maximum number of points in the training partition. We fit the final model for each CV partition using the combination of power parameter and number of observations that resulted in the lowest three-fold cross-validated RMSE on data in the training partition.

*Investigating possible overfitting*

After models were fitted, we looked for evidence of overfitting by inspecting graphs of 1) the number of predictor variables used by GLMs as a function of sample size, and 2) prediction performance (spatial block cross-validated AUC) as a function of the number of terms in GLMs and sample size. We also explored the effect of the constraints we placed on the computation time of boosted regression trees (i.e. limiting models to 30,000 or fewer trees) by inspecting boxplots of the number of trees used to fit models as a function of sample size and spatial sampling bias. Finally, we assessed the effect of species prevalence on model performance metrics by inspecting plots of AUC and RMSE as a function of species prevalence and as a function of the number of positive detections of the focal species in the test dataset.

**Analyzing effects of sampling bias and sample size**

Our main analysis (reported in the main text) used boosted regression trees to model the predictive performance (AUC and RMSE) of SDMs as a function of spatial sampling bias and sample size (average number of observations per species), SDM method, and (in the case of RMSE) species prevalence. To assess whether our conclusions depended on the modeling method, we also used GAMs (Wood, 2017) to perform the same analysis of AUC and RMSE of SDMs as a function of spatial sampling bias and sample size (average number of observations per species), SDM method, and species prevalence. Using both boosted regression trees and GAMs provided a simple sensitivity test to ensure that our conclusions were not dependent on the choice of modelling method. We also used the GAM predictions of AUC to produce the contour lines in Fig. 8 of the main text because the smoother GAM function made the shape of the contour lines easier to visually distinguish than contour lines produced with boosted regression tree predictions.

We used boosted regression trees to model AUC as a function of a categorical spatial sampling bias variable, the average number of observations per species, and SDM method. Boosted regression trees used a tree complexity of 3, a learning rate of 0.001, a Gaussian distribution, and the number of trees selected by the 'gbm.step' function in the 'dismo' R package.

We fit GAMs to model AUC and RMSE using the 'gam' function in the 'mgcv' R package. We fit separate GAMs to model the prediction performance of each of the four SDM modelling methods because three-way interactions cannot be specified in 'gam' and we expected three-way interactions. We modeled AUC as a function of a categorical spatial sampling bias variable and a smooth of the average number of observations per species by sampling bias (so that the response shape of AUC to sample size could vary with bias level). We modeled RMSE as a function of a categorical spatial sampling bias variable, a smooth of the average number of observations per species by sampling bias, and a smooth of species prevalence. We used a beta distribution and logit link, and smoothed the number of observations per species by sampling bias level using cubic regression splines with a basis dimension of five. The basis dimension was chosen by fitting multiple models with basis dimensions varying from two to six and looking at effective degrees of freedom and the shape of fitted smooths. We selected the basis dimension to be high enough that effective degrees of freedom were below the basis dimension and neither the shape of the smooth nor the basis dimension changed substantially when the basis dimension was increased.

Predictions were generated from fitted boosted regression trees and GAMs. We compared the expected value of AUC and RMSE for SDMs trained with data containing different amounts of spatial sampling bias and different sample sizes. Variable importance was assessed based on the reduction in squared error attributed to each variable in the boosted regression tree models and based on the change in adjusted $R^2$ of GAMs when variables were removed from the full model.

### Supplementary Results & Discussion

#### Prediction performance of SDMs

AUC was lower when models were evaluated using spatial block cross-validation than when models were evaluated on the training data, as expected (Fig. S9). Model evaluation on training data is known to be overly optimistic in most cases (Roberts et al., 2017), and our results confirm that.

Analyses of the effects on prediction performance of spatial bias, average number of records per species, and SDM method were qualitatively similar when analyzed using boosted regression trees and GAMs, suggesting that our conclusions did not depend on the choice of error distribution or modeling technique (Fig. 5, Fig. S9).

*Investigating possible overfitting in GLMs and the effects of limiting computation time for boosted regression trees*

To avoid overfitting the GLM species distribution models, we used fewer terms in models when sample size was small, and only allowed more terms when sample size was large (Fig. S10). The poor performance of GLM species distribution models trained with small sample sizes cannot be attributed to overfitting, as GLMs used relatively few terms when sample size was small (Fig. S10). Out-of-sample prediction performance (AUC) of GLMs increased with the number of terms in models up to about three or four terms (Fig. S11). When sample size was intermediate (an average of 10 or 50 records per species), prediction performance then initially increased with the number of terms in models, then decreased when more terms were used, indicating possible overfitting (Fig. S11, panels C, D, and E). The possible overfitting was most pronounced for models trained with median or severely biased data with an average of 50 records per species (Fig. S11, panel D). This suggests that GLMs may have been overfitting at moderate sample sizes, despite us limiting the number of terms in models based on sample size. There was no evidence of overfitting at small sample sizes, mainly because models were restricted to using very few predictor variables (Fig. S11, panels A and B). Prediction performance of GLMs generally increased with sample size (Fig 5 in the main text), despite the evidence of possible overfitting at intermediate sample sizes suggested by Fig. S11. More careful model selection and control of overfitting in GLMs, for example by selecting the final model terms using cross-validation, could increase the prediction performance of models trained with moderate sample sizes even further. However, this would not change our main findings, and in fact would strengthen the pattern of prediction performance increasing with sample size (Fig. 5). Therefore, we do not think the evidence of some overfitting in GLMs affects the main conclusions of this study, namely that prediction performance of species distribution models is affected more strongly by sample size and species distribution modelling method than by spatial sampling bias.

We limited boosted regression tree species distribution models to using fewer than 30,000 trees. However, most models fit with fewer than 10,000 trees (Fig. S12), and prediction performance was unrelated to the number of trees used, as long as number of trees was above about 2,000 (Fig. S13). Boosted regression trees failed to fit models for some species, especially when sample sizes were small (Fig. 4 of main text), perhaps because we abandoned model fitting if models did not successfully fit with 30,000 or fewer trees. It is possible that given more computation time and larger numbers of trees, boosted regression trees could successfully fit models to more species. However, our assessment of the prediction performance of models was based only on models that *did* successfully fit. For those models that fit, we saw no indication that the prediction performance was limited by permitting models to use only up to 30,000 trees (Fig. S13). Rather, the majority of models fit with far fewer than 30,000 trees, (Fig. S12), and prediction performance was generally constant for models with numbers of trees from about 2,000 to 30,000 (Fig. S13). Any practical species distribution modelling will be done within the constraints of available computational resources. We do not think that increasing the maximum permissible number of trees for boosted regression trees above 30,000 would change the main conclusions of this study, namely that prediction performance of species distribution models is affected more strongly by sample size and species distribution modelling method than by spatial sampling bias.

*Small community simulation*

Results from the small community simulation were qualitatively similar to results from the large community simulation (Fig. S14). Prediction performance was similar when models were trained with data showing no spatial bias or low spatial bias. Prediction performance was lower when models were trained with data with median or severe spatial bias, at least for GLMs (Fig. S14). In the small community simulation, inverse distance-weighted interpolation appeared to be less affected by spatial bias than it was in the large community simulation. Despite this, GLMs trained with severely spatially biased data still outperformed the best inverse distance-weighted interpolation models (Fig. S14).
