## Supplemental Figure S3 for "Data quantity is more important than its spatial bias for predictive species distribution modelling"

A

| checklist ID | longitude | latitude | list length | species name |
|---|---|---|---|---|
| L14 | 75000 | 85000 | 4 | sp2 |
| L14 | 75000 | 85000 | 4 | sp6 |
| L14 | 75000 | 85000 | 4 | sp5 |
| L14 | 75000 | 85000 | 4 | sp1 |

B

| checklist ID | longitude | latitude | list length | sp 1 | sp 2 | sp 3 | sp 4 | sp 5 | sp 6 | sp 7 | ... | sp240 |
|---|---|---|---|---|---|---|---|---|---|---|---|---|
| L14 | 75000 | 85000 | 4 | 1 | 1 | 0 | 0 | 1 | 1 | 0 | ... | 0 |
