## Supplemental Table S7 for "Data quantity is more important than its spatial bias for predictive species distribution modelling"

**Table S7. Adjusted R^2^ of generalized additive models (GAMs) modeling prediction performance (AUC) of species distribution models for simulated species.** The full model (1^st^ line) models AUC as a function of spatial bias and a smooth of sample size by spatial bias. The 2^nd^ and 3^rd^ lines show the the adjusted R^2^ of models with the sample size or spatial bias terms removed, respectively. Removing the sample size term reduces the adjusted R^2^ more than removing the spatial bias term does.

| **Model** | **Adj. R^2^ (GLM SDMs)** | **Adj. R^2^ (inverse distance-weighted interpolation SDMs)** | **Adj. R^2^ (boosted regression tree SDMs)** |
| --- | --- | --- | --- |
| full model | 0.45 | 0.22 | 0.13 |
| - spatial bias | 0.43 | 0.21 | 0.11 |
| - sample size | 0.01 | 0.01 | 0.03 |
