## Supplemental Table S8 for "Data quantity is more important than its spatial bias for predictive species distribution modelling"

**Table S8. Relative influence of variables on the root mean squared error of species distribution models of simulated species (large community simulation).** The relative influence for each variable is the reduction in squared error attributable to that variable in a boosted regression tree model.

| Variable | Relative importance (reduction in squared error) |
| --- | --- |
| Species prevalence | 99.85 |
| Average number of records per species | 0.1494 |
| Species distribution modelling method | 0.0032 |
| Spatial bias | 0.0001 |
